# YAP1 and QSER1 are Key Modulators of Embryonic Signaling Pathways in the Mammalian Epiblast

**DOI:** 10.1101/2025.06.16.659935

**Authors:** Elizabeth Abraham, Thomas Roule, Aidan Douglas, Emily Megill, Olivia M. Pericak, Jordan E. Howe, Carmen Choya-Foces, Joanne F. Garbincius, Henry M. Cohen, Paula Roig-Flórez, Mikel Zubillaga, Mark D. Andrake, Seonhee Kim, John W. Elrod, Naiara Akizu, Conchi Estaras

## Abstract

YAP1 signaling is essential for development but its specific roles in early embryogenesis remain poorly understood. To shed light on this, we analyzed YAP1’s role in regulating the pluripotency of the mammalian epiblast, using scRNAseq approaches. Conditional deletion of *Yap1* in the mouse epiblast (*Sox2*-Cre) altered the expression of signaling genes, including *Nodal*, *Wnt3*, and *Fgf8*. Accordingly, *Yap1* loss led to enhanced differentiation of the epiblast toward primitive streak lineages, as evidenced by the upregulation of T/*Brachyury* and *Eomes* genes. Furthermore, a proximity labeling assay in human pluripotent stem cells, followed by biochemical assays and molecular modeling predictions, revealed that YAP1 cooperates with QSER1 protein to regulate lineage genes. Our analysis shows that YAP1:TEAD4 enhancers recruit QSER1 to prevent RNA Polymerase II recruitment. Accordingly, QSER1 depletion, similar to YAP1, increases NODAL gene expression and leads to hyperactive NODAL signaling in human 2D-gastruloids. Overall, our findings define a role of YAP1 in the epiblast in vivo and uncovered an interplay with QSER1 controlling the activity of developmental signaling pathways in pluripotent cells.

## INTRODUCTION

The main transcriptional effectors of the Hippo signaling pathway are YAP1 and its homolog TAZ. When YAP1 and TAZ are phosphorylated by Hippo kinases, they are degraded in the cytoplasm. When dephosphorylated, YAP1 and TAZ are stabilized and translocated into the nucleus where they interact with TEAD1-4 DNA binding proteins to induce the expression of genes that promote proliferation and growth^1^. Thus, loss-of-function mutations in the core kinases or overexpression of YAP1 and TAZ up-regulates the expression of genes that induce cell proliferation, cell survival, and tissue growth. Notably, TAZ knockout (KO) mice complete embryonic development, whereas YAP1 KO results in early embryonic lethality, underscoring a more prominent role of YAP1 in embryogenesis that TAZ cannot compensate^2,3^. Overall, the results of these experimental approaches established the Hippo:YAP1 signaling pathway as a universal regulator of organ size^4^. However, emerging lines of research underscored that the physiological roles of YAP1 may be unrelated to cell proliferation or growth in some developmental contexts. During liver development, YAP1 overexpression causes overgrowth^5^, but conditional deletion of YAP1/TAZ in the liver lineage does not prevent this organ from reaching its normal size^6,7^. Instead, endogenous YAP1 activity is required for cholangiocyte differentiation^6^. Similarly, *Yap1* KO mouse embryos can initiate gastrulation and develop until ∼E8 without noticeable changes in size, suggesting that YAP1 is not required for embryonic growth or proliferation of totipotent, multipotent, and pluripotent stem cells^3^. Instead, during this embryonic period, YAP1 regulates zygotic genome activation in totipotent cells and specification of trophectodermal cells in multipotent cells^8,9^. However, whether YAP1 regulates the differentiation of pluripotent stem cells in vivo, is still unknown.

The natural progression of the pluripotent cells of the epiblast, triggers gastrulation^10^. During gastrulation, the pluripotent epiblast differentiates into each of the three germ-layer lineages, breaking the symmetry of the embryo and establishing the body plan. The formation of the primitive streak (PS) in the posterior part of the epiblasts marks the beginning of gastrulation. It gives rise to mesoderm and endoderm layers, while the anterior epiblast commits to ectodermal lineages^10–12^. The activity of key signaling pathways, including WNT, NODAL and BMP, enables selective activation of gene-expression programs necessary for germ-layer specification^13^. Before gastrulation, NODAL signaling maintains pluripotency of the epiblast^14–17^ while WNT signaling is initially low^18,19^. Upon the onset of gastrulation, upstream signals from extraembryonic tissues— including BMP4—stimulate *Wnt3* expression in adjacent epiblast cells^12^. WNT signaling activation stabilizes β-CATENIN, which partners with Nodal transcriptional effectors SMAD2/3 to drive PS gene expression, including *Eomes* and *T*/*Brachyury*, in the posterior part of the epiblast^20,21,22^. Conversely, in the anterior epiblast, WNT and NODAL activity are actively suppressed to permit ectodermal differentiation, an inhibition largely mediated by WNT and NODAL antagonists secreted by extraembryonic tissues^13,23,24^. Thus, as development progresses, NODAL signaling becomes progressively confined to the posterior epiblast to induce the PS in concert with WNT^20,21,22^, enabling ectoderm commitment in the anterior region. Overall, precise regulation of WNT and NODAL signaling is necessary to ensure correct lineage differentiation and anterior-posterior patterning during gastrulation.

Using human pluripotent stem cell (hPSC) models, we and others have shown that YAP1 regulates germ-layer differentiation by repressing the activity of WNT and NODAL signaling^25–27^. Furthermore, our recent studies revealed that *Yap1-*depleted embryos have altered proportion of germ-layer derivatives, compared to controls^28^, suggesting that *Yap1* loss disrupts epiblast differentiation in vivo. Thus, in the present study we investigate the molecular mechanisms by which YAP1 regulates cell-fate decisions of the pluripotent epiblast. Single-cell transcriptomic profiling of embryos at E7 revealed that conditional YAP1 deletion led to upregulation of key gastrulation signaling genes, including *Nodal*, *Wnt3*, and *Fgf8* in the epiblast. Furthermore, E7 mutant embryos display an increased proportion of cells in the PS, compared to controls, and increased expression of PS genes, including *T/Bra* and *Eomes*. We then conducted a proximity-based proteomic screen in hESCs, which identified QSER1 as a high-confidence nuclear interactor. Accordingly, by ChIP-seq analysis we found that QSER1 co-localizes with YAP1 and TEAD4 on a subset of developmental enhancers, including those regulating *NODAL* and *SMAD2*. Functional and structural analyses revealed that YAP1 serves as a bridge between QSER1 and TEAD4, facilitating QSER1 recruitment and stabilizing a TEAD-YAP1-QSER1 trimeric complex in the chromatin. Finally, we found that loss of QSER1 on YAP1:TEAD4 enhancers increased RNA polymerase II occupancy on these sites and phenocopies YAP inhibition, leading to ectopic NODAL expression and mesoderm expansion in hESC-derived 2D gastruloids. Altogether, our findings define a role in YAP regulating epiblast differentiation in vivo and reveal a previously unrecognized YAP1–QSER1 module that functions to fine-tune gene expression during lineage differentiation.

## RESULTS

### Conditional deletion of *Yap1* (*Yap1* cKO) in the epiblast and analysis of E7 embryos

To investigate the functional role of YAP1 in epiblast differentiation *in vivo*, we conditionally deleted *Yap1* in mouse embryos using a Sox2-Cre driver^29^. *Sox2* gene expression initiates at ∼E5.5 in the epiblast, and *Cre*-mediated recombination occurs throughout the epiblast and its derivatives, excluding earlier arising extraembryonic lineages^29^. We performed single-cell RNA-seq analysis on freshly isolated embryos at E7 prior to genotyping of Cre, flox, and sex alleles, using a methodology optimized in our lab^30^ (**Figure EV1A-D**). We sequenced a pool of three *Yap1* cKO (*Yap1*^Flox/Flox^: Sox-Cre) and four heterozygous control littermates (*Yap1*^Flox/+^: Sox2-Cre) (**Figure 1A and Figure EV1C).** A total of 1,829 control cells and 1,897 *Yap1* cKO cells were initially sequenced. After excluding cells that did not meet specific data quality standards, these numbers were reduced to 788 and 862, respectively (see **Methods** for details).

**Figure 1.**
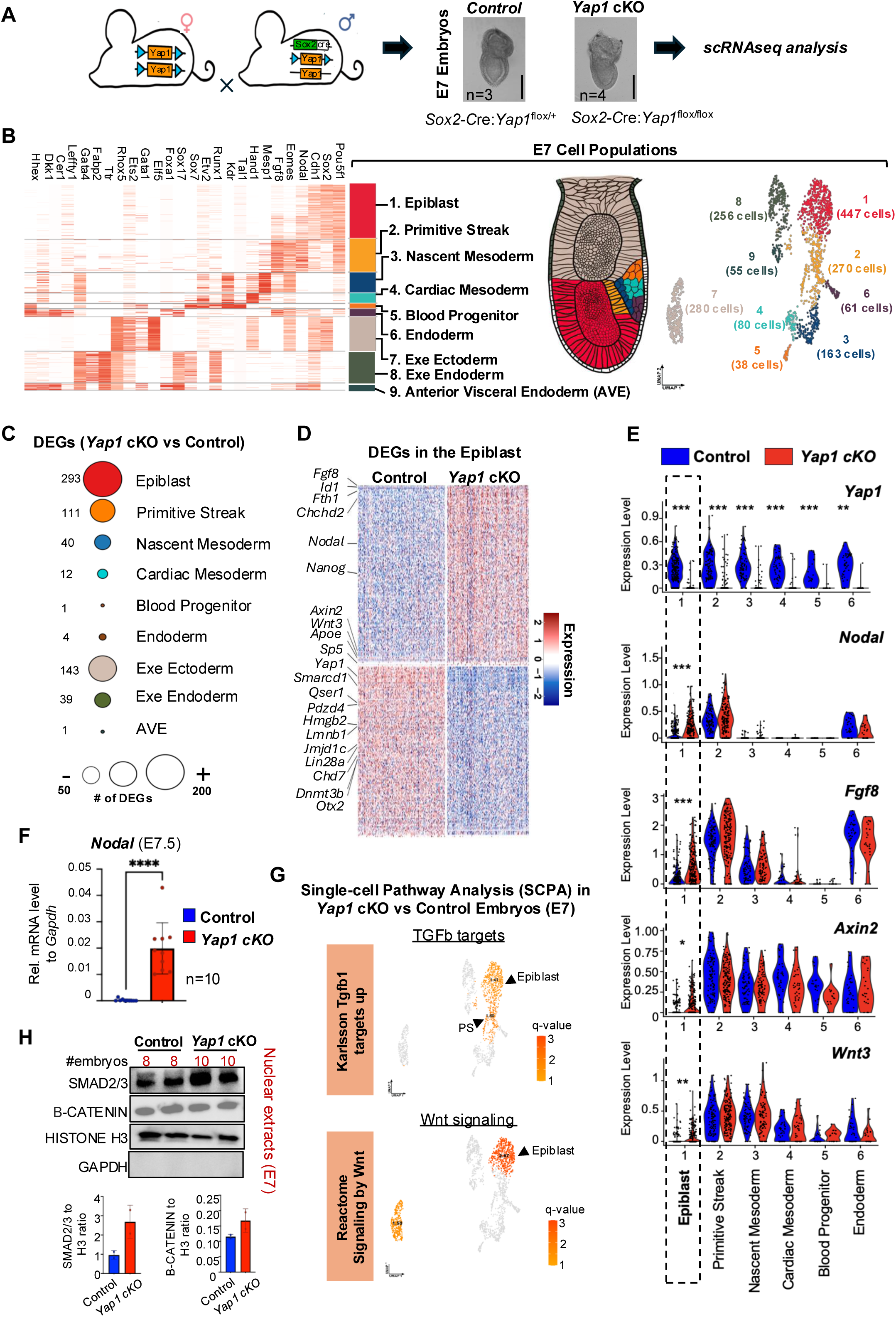
Nodal and Wnt signaling genes are activated in the epiblast of *Yap1 cKO* embryos compared to controls. (A) Mice scheme shows the breeding strategy to obtain embryos with conditional deletion of *Yap1* in the epiblast (Sox2cre). Blue arrowheads indicate LoxP alleles. E7 heterozygous control (Sox2cre:YAPflox/+) and *Yap1* cKO (Sox2cre:Yap1flox/flox) embryos were processed for scRNAseq analysis. Bright-field images show representative embryos of the indicated genotype. The number of embryos processed for sequencing is indicated. Scale Bar 250µm. (B) Heatmap showing expression of lineage markers used to annotate cell populations in the E7 scRNAseq datasets. On the right, a schematic of an E7 mouse gastrula and a UMAP of E7 scRNAseq showing the detected cell populations with the number of cells in parentheses, color-coded to match the heatmap. (C) Dot plot depicts the number of differentially expressed genes (DEGs) in each cluster, with the exact count indicated to the left of each dot. Note that the epiblast cluster contains the highest number of DEGs (abs(Log2FC)>0.25, adj. p-value <0.05). (D) Heatmap shows DEGs in the epiblast of *Yap1* cKO versus control embryos. Relevant genes for pluripotency and differentiation are shown. (E) Violin plots show scRNAseq expression levels of indicated genes across clusters in control and *Yap1* cKO. The dotted box highlights the epiblast cluster. (Adjusted p-value is based on Bonferroni correction, ^∗^*p* < 0.05, ^∗∗^*p* < 0.001, ^∗∗∗^*p* < 0.0001.) (F) Graph shows RT-qPCR analysis of the *Nodal* gene in E7.5 control and *Yap1* cKO embryos. (n=10) Data presented as mean ± SEM. Statistical analysis: Student’s t-test, ****p<0.0004 (G) Single-cell pathway enrichment analysis (SCPA) was performed on DEGs of *Yap1* cKO compared to control. The UMAP plot shows the enrichment of two of terms related to the Nodal/TGFb and Wnt pathway. Significant q-values (>1.4) are displayed in orange with the names of populations. The complete list of Wnt and TGFb terms enriched are shown in **Figure EV1I**. (H) Western blot of nuclear extracts of E7 control and *Yap1* cKO embryos. The number of embryos pooled per lane is indicated above each lane, along with the markers analyzed. Uncropped blots are found in **Figure EV1J**.

Sequenced cells were manually annotated based on the expression of known markers^31–33^. We identified six embryonic (epiblast, primitive streak (PS), nascent mesoderm, cardiac mesoderm, blood progenitor, endoderm) and three extraembryonic populations (extraembryonic endoderm, anterior visceral endoderm (AVE), and extraembryonic ectoderm) in the E7 embryos, based on the expression of known markers (**Figure 1B** and see **Methods** for details). Importantly, *Yap1* cKO embryos showed a significant reduction in Yap1 mRNA levels across embryonic clusters compared to controls **(Figure 1E** and **Figure EV1E)**. Consistent with reduced YAP1 activity in the embryos, we observed typical YAP1 target genes and Hippo regulators transcriptionally affected. These include *Ccn2* (also known as *Ctgf*), which was significantly downregulated in the primitive streak and nascent mesoderm, and *Ccn1* (also known as *Cyr61*), which was reduced in the epiblast. Regulators of the Hippo pathway, including *Amotl2*^34^, *Ptpn14*^35^, and *Wwc2*^36^, were also downregulated across multiple lineages, such as the epiblast, primitive streak, and cardiac mesoderm **(Table EV1 and Figure EV1F)**. These results support the effective disruption of YAP1 activity in the cKO embryo. Importantly, the expression of the *Wwtr1* gene, which codifies for the YAP1 homolog TAZ, was not significantly affected by *Yap1* deletion (**Figure EV1E**). As anticipated, *Yap1* expression in extraembryonic clusters remained similar between *Yap1* cKO embryos and controls (**Figure EV1E**). Cell cycle analysis revealed no significant differences in the *Yap1* cKO compared to control embryos (**Figure EV1G**). In agreement, control and *Yap1* cKO E7 embryos display similar cell count and epiblast perimeter (**Figure EV1H**). We conclude that our experimental approach effectively captured the major cell populations of E7 embryos, achieved efficient conditional *Yap1* deletion, and showed that *Yap1* loss did not alter cell cycle dynamics compared to controls.

### YAP1 represses Nodal and Wnt signaling pathway genes in the epiblast

Differential expressed genes (DEGs) analysis between *Yap1* cKO and control embryos revealed varying numbers of genes impacted by *Yap1* deletion across embryonic clusters (**Figure 1C and Table EV1**). Within the embryonic populations, the epiblast cluster exhibited the highest number of DEGs (293; 151 upregulated, 142 downregulated, adj. p <0.05), followed by the PS (111) and the nascent mesoderm (40) (**Figure 1C**). These findings align with previous studies showing a significant role of YAP1 in the transcriptome of pluripotent stem cells in vitro^21,26,27,37^. Among the downregulated genes in the epiblast, we identified key pluripotent regulators such as *Jmjd1c*^38^, *Dnmt3b*^39^, and *Qser1*^40^. Conversely, upregulated genes included the critical signaling pathway genes *Nodal*, *Wnt3,* and *Fgf8* (**Figure 1D-F**). Along with Wnt and Nodal, Fgf signaling also promotes epiblast differentiation toward PS cell-fates^18,41–43^. Consistent with this, we observed upregulation of downstream targets of these pathways, including *Axin2*, *Id1*, and *Nanog*. *Axin2* is a typical β-CATENIN target gene, thought to fine-tune Wnt signaling activity in the epiblast^44^. *Nanog* and *Id1* are direct targets of SMAD2/3 involved in the repressing of neuroectodermal fates^17,45,46^ and regulating epiblast differentiation timing and cardiac mesoderm differentiation^47–49^, respectively. Furthermore, single-cell Pathway Analysis (SCPA) retrieved TGFβ/NODAL and WNT signaling related terms specifically enriched in the epiblast cluster of the *Yap1* cKO embryos, compared to controls (**Figure 1G and EV1I**). Accordingly, we detected an accumulation of nuclear SMAD2/3, the transcriptional effectors of Nodal signaling, in nuclear extracts of mutant embryos, compared to controls, further suggesting an increase in the activity of the NODAL pathway upon *Yap1* loss (**Figure 1H and Figure EV1J (nuclear) and Figure EV1K (total extracts)**). However, we failed to detect nuclear accumulation of the WNT effector β-CATENIN in the mutant embryos, compared to controls (**Figure 1H and Figure EV1J (nuclear) and Figure EV1K (total extracts)**), which could be due to a milder activation of the pathway combined with the dynamic nature of β-CATENIN nuclear localization, as previously noted by others^50^.

Overall, these data show that conditional deletion of *Yap1* in the epiblast affects the expression of genes important for pluripotency and differentiation. Furthermore, we conclude that the predominant role of YAP1 in the epiblast is to repress the expression of signaling genes regulating PS differentiation.

### Conditional YAP1 deletion leads to an expanded Primitive Streak (PS) domain at E7

To test how YAP1 deletion in the epiblast affects E7 populations, we analyzed the proportion of cells in each cluster in *Yap1* cKO embryos and controls. Among the embryonic clusters, the PS was the only cluster with significant changes. *Yap1* cKO embryos displayed a significant increase in the number of PS cells, compared to controls (**Figure 2A).** Two of the extraembryonic clusters also displayed significant differences in the number of cells (**Figure EV2A).** However, the extraembryonic clusters do not express the *cre* allele and preserve YAP1 expression (see **Figure EV1E**). Therefore these differences may be due to a non-autonomous effect of *Yap1* deletion in the epiblast.

**Figure 2.**
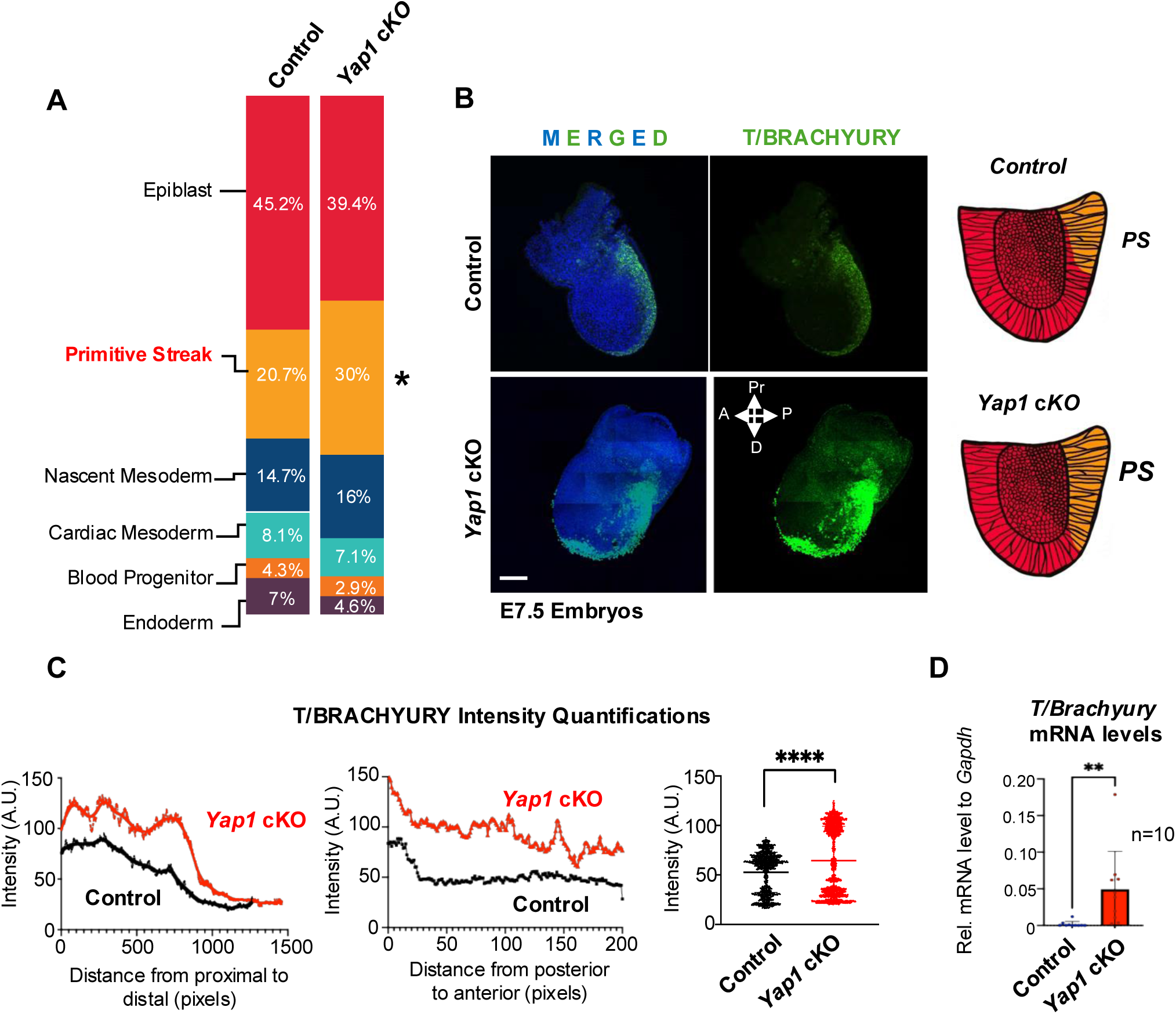
Yap1 loss in the epiblast increases differentiation toward the primitive streak (PS). (A) Bar graph showing the percentage of embryonic cell populations detected by scRNA- seq analysis in control and *Yap1* cKO embryos. Statistical significance was assessed using the Chi-square test (*p < 0.05). Only embryonic populations are shown. See Figure EV2A for analysis including all populations. (B) Representative images of whole-mount immunostaining for the PS marker BRACHYURY (T/BRA) (green) in E7.5 control and *Yap1* cKO embryos. DAPI (blue) marks nuclei. On the right, a scheme summarizing results; compared to controls, *Yap1* cKO embryos have expanded the PS domain. Scale Bar 250µm Pr: proximal, A: anterior, P: posterior, D: Distal. (C) Graphs show quantifications of T/BRA signal intensity along the proximal to distal axis of the embryo (left), the posterior to anterior axis (middle), and the overall intensity of the immunostaining (right). An in-house developed Matlab script was applied to quantify fluorescence. The experiment was replicated with three separate litters. (n=3) Data are presented as mean ± SEM. Statistical analysis: Student’s t-test, ****p<0.0004. (D) RT-qPCR of *T/Bra* in E7.5 control and *Yap1* cKO embryos. (n=10) Data are presented as mean ± SEM. Statistical analysis: Student’s t-test, ^∗∗^*p* < 0.001.

To visualize the PS domain in gastrulating embryos, we analyzed the expression pattern of *T/Brachyury*, a general marker of the PS^51–53^, using whole-mount immunostaining. At the onset of gastrulation, *T/Brachyury* is initially expressed in the proximal posterior epiblast and expands both anteriorly and distally as development progresses^52^. In *Yap1* cKO embryos, we observed a significant increase in *T/Brachyury* expression levels, along with a distal expansion of its expression domain compared to control embryos (**Figure 2B-D and Figure EV2B**). Therefore, we conclude that YAP1 deletion leads to enhanced PS differentiation of the epiblast.

### YAP1 deficiency disrupts embryo patterning

Premature or excessive differentiation toward PS may result in severe consequences for gastrulation patterning. For instance, it may lead to an imbalance in the proportion of populations derived from the PS and/or a depletion in the pool of epiblast cells left for ectodermal commitment.

Indeed, older *Yap1* cKO embryos display severely underdeveloped headfolds and neural plate structures (**Figure EV2C-D**) that are not compatible with life passed E8.5^3, 28^. These abnormalities are consistent with impaired ectoderm development and disruption of anterior structures^57^.

To further determine the developmental consequence of epiblast Yap1 deletion, we analyzed the proportion of cell populations in the scRNAseq of E7.75 embryos we previously generated^28^. Remarkably, a comparison of epiblast-derived populations between control and *Yap1* cKO at E7.75 revealed an increase in the proportion of hematopoietic stem cell and endoderm populations alongside a significant reduction in neural and surface ectoderm populations in *Yap1* cKO embryos (**Figure EV2E**). Other mid-posterior PS populations, including presomitic and somitic mesoderm, were also underrepresented in the *Yap1* cKO embryos (**Figure EV2E**). To confirm these results, we performed qPCR and Western blot analyses of the anterior primitive streak marker *Eomes*^54^, endoderm marker *Foxa2*^55^ and neuroectoderm marker *Otx2*^56^ in E7.75 embryos. Consistent with the morphological analysis and scRNAseq data, *Eomes* and *Foxa2* were upregulated and *Otx2* downregulated in *Yap1* cKO (**Figure EV2F-G**).

Overall, we conclude that conditional *Yap1* deletion leads to imbalanced differentiation of the epiblast. Specifically, *Yap1* loss in the epiblast promotes the expansion of anterior primitive streak derivatives while compromising neural committed cells. This shift is consistent with increased NODAL and WNT signaling activities in the mutant embryos, as suggested by our transcriptomic analysis of the epiblast at E7.

### Analysis of YAP1 nuclear interactors in human pluripotent stem cells

In many cell contexts, YAP1 regulates transcription of genes controlling proliferation and cell growth, such as *Myc* or *Ctgf*^58^. However, in the mouse epiblast (**Figure 1**) and in hESCs^25,37^, YAP1 also regulates genes linked to differentiation, like *Wnt3* and *Nodal*. This suggests the existence of a pluripotent-specific YAP1-transcriptional network. To gain insight into this, we performed a proximity biotinylation assay to identify YAP1-associated nuclear proteins in hESCs. Although key differences exist between mouse epiblast and human hESCs^10,59,60^, they share important functional similarities, including their pluripotent state and capacity to give rise to all three germ layers^61,62^. Accordingly, the phenotype of YAP1 deletion in hESCs^25^ closely mirrors the phenotype of *Yap1* cKO in the epiblast (**Figure 1** and **Figure 2**).

We carried out a proximity biotinylation (BioID2^63^) assay on nuclear extracts of hESCs using YAP1 as bait (**Figure 3A**). For this, we created a inducible hESC clonal line that expresses YAP1 fused to Bir2 under the control of a Doxycycline responsive promoter. Doxycycline and biotin concentrations were optimized to observe a stable nuclear YAP1-MYC-BIR2 expression (**Figure 3B and Figure EV3A**) and effective biotinylation of nuclear proteins, respectively (**Figure 3C and Figure EV3B**). Furthermore, we confirmed that the YAP1-MYC-BIR2 fusion protein retains the ability to interact with TEAD4, by co-immunoprecipitation experiments (**Figure EV3C**) and ChIP-qPCR approaches (**Figure EV3D**), to further confirm the functionality of our YAP1-MYC- BIR2 construct.

**Figure 3.**
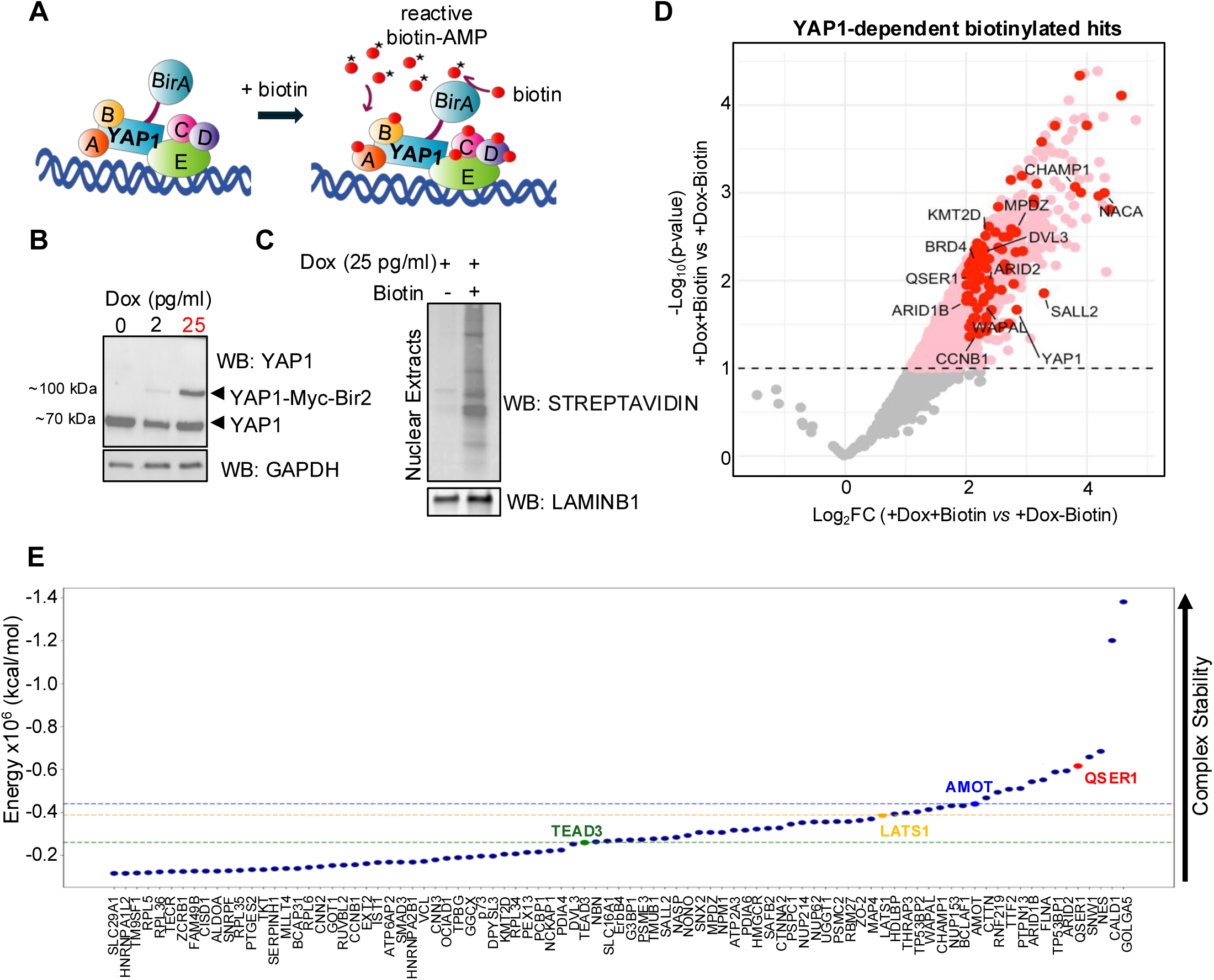
BioID2 assay identifies QSER1 in the proximity network of YAP1 in hESCs (A) Scheme of BioID2 assay to identify potential nuclear interactors of YAP1 in hESCs. (B) A doxycycline-inducible YAP1-Myc-Bir2 clonal cell line was generated in H1 hESCs. Western blot of YAP1 shows expression of construct following the indicated treatments. 25pg/ml of Doxycycline (Dox) was used for experiments (highlighted in red). Note that the expression of the YAP1-Myc-Bir2 construct at 25pg/ml (∼90KDa band) is similar to the expression of endogenous YAP1 (∼70KDa band). GAPDH was used as a loading control. (C) Western blot of YAP1-Myc-Bir2 against Streptavidin shows biotinylated nuclear proteins in the presence and absence of biotin, as indicated. LAMINB1 was used as a loading control of nuclear extracts. (D) Volcano plot showing protein hits detected in the BioID2 assay. Gray dots indicate proteins not significantly enriched in +Biotin+Dox vs. –Biotin+Dox condition. Pink dots represent proteins significantly enriched (Log2FC > 2 and adjusted p-value < 0.05) in the +Biotin+Dox vs. –Biotin+Dox comparison, totaling 311 hits. A secondary filter was applied to the 311 hits: red dots highlight proteins also enriched (Log2FC > 1.5) in +Biotin+Dox compared to both +Biotin–Dox and –Biotin–Dox conditions. Only proteins with LFQ values in at least two of three replicates were included in the analysis. The name of some hits is shown; the full list is provided in **E.** (E) Computational protein docking analysis of YAP1 with 83 candidate interactors identified by BioID2. The plot displays the predicted potential energies of YAP1 complexes with each candidate, which serve as a proxy for the stability of the protein complex. Energies for three known YAP1 interactors (TEAD3, LATS1, and AMOT) are included as positive controls. The QSER1–YAP1 complex is highlighted and ranks as the fifth most stable predicted interaction.

Mass spectrometry analysis of purified streptavidin-bound peptides was performed in four experimental conditions, as shown in **Figure EV3E**. The analysis of the potential hits was performed as follows. We first compared biotin-treated YAP1-MYC-BIR2 samples (+dox,+biotin) to untreated YAP1-MYC-BIR2 samples (+dox,+biotin) and found 311 hits significantly enriched (n=3, Log2FC>2, FDR<10%) (**Figure 3D**, highlighted as pink dots in volcano plot). Then, we filtered out these 311 hits list, by discarding the proteins that were still biotinylated in the absence of YAP1-MYC-BIR2 construct (-dox, +biotin) or the absence of YAP1-MYC-BIR2 construct and biotin (-dox, -biotin) (See Methods for details). After these filters, we obtained a subset of 83 proteins (**Figure 3D**, highlighted as red dots in volcano plot), as bonafide biotinylated targets of YAP1.

Among the 83 biotinylated hits, we found transcriptional co-activators and co-repressors reported to interact with YAP1, including the epigenetic reader BRD4^64^, the subunit of the chromatin remodeler complex SWI/SNF ARID2^65^, and the methyltransferase KMT2D^66^. Additionally, we detected other well-known interactors of YAP1, such as the subunit of the Crumbs cell polarity complex MPDZ^67^, and DVL3, a core component of the Wnt signaling pathway, that shuttles between the nucleus and cytoplasm and regulates YAP1’s intracellular location^68^ (**Table EV2 and Figure 3D-E**). The analysis also retrieved new potential interactors, including the zing finger protein SALL2^69^, the cohesin regulatory protein WAPAL^70^, or the splicing and transcription regulator SNW1^71^. QSER1, a recently characterized protein, whose activity has been tightly associated with the protection of DNA methylation sites in hESCs was also among the 83 gene list^40,72^ (**Figure 3D-E** and **Table EV2**). Nonetheless, our mass spectrometry analysis detected few TEAD peptides, which did not reach the threshold for significant enrichment (see raw values in **Table EV2** and **Figure EV3E**). Other published BioID/TurboID datasets report similarly low TEAD recovery when YAP1 was used as bait^73^, which could be due to limited accessible lysines, rigid structure of TEAD or/and geometric constraints due to the tight YAP1:TEAD interactions^73–75^.

### Computational Docking Predicts High Stability for a QSER1:YAP1 Complex

To prioritize potential YAP1 interactors among the hits identified in the BioID2 assay for downstream analysis, we performed computational docking to evaluate whether these proteins could plausibly form stable complexes with YAP1. Structural models from AlphaFold^1–3^ were used to simulate binding interactions using HDock^4–8^, and each complex was assessed for stability through energy minimization^11^. This analysis allowed us to rank candidate interactors based on predicted structural compatibility, with lower energies associated to more stable complexes (see methods for details). As a reference, we included known YAP1 interactors—TEAD3^76^, LATS1^77^, and AMOT^78^—to interpolate benchmark energy values (**Figure 3E**). Among these positive controls, the AMOT:YAP1 complex exhibited the lowest energy. Notably, 10 of the 83 candidate proteins displayed even lower predicted energies than AMOT, including transcriptional regulators such as ARID1B and SNW1. Among these, the QSER1:YAP1 complex ranked within the top five most stable predictions. Considering its predicted binding stability along with QSER1’s prominent yet poorly understood role in pluripotency^40^ we chose to investigate this potential functional interaction in greater detail. To validate the predicted interaction between QSER1 and YAP1, we performed co-immunoprecipitation (Co-IP) assays in the YAP1-MYC-BIR2 hESCs lines, which confirmed that YAP1 interacts with endogenous QSER1 (**Figure EV3C**). Immunofluorescence analysis further revealed that both proteins co-localize in the nucleus of hESCs (**Figure EV3F-H**). In addition, we conducted size-exclusion chromatography experiments using nuclear extracts of hESCs, which showed that QSER1 and YAP1 co-elute in overlapping fractions of high molecular weight, suggesting they are part of same nuclear complexes (**Figure EV3I**). Together, these results provide strong evidence that QSER1 and YAP1 interact within the nucleus of hESCs.

### QSER1 is an intrinsically disordered protein enriched at promoters and enhancers in hESCs

Previous finding identified QSER1 through a methylation-based screen as a TET-interacting partner in hESCs^40^ and found a role for QSER1 in protecting DNA methylation, particularly at EZH2-bound loci^40^. However, ChIP-seq analyses of a FLAG-tagged QSER1 protein in hESCs revealed broad binding to enhancer regions, beyond EZH2 co-occupancy^40^, suggesting additional regulatory roles yet to be defined.

To further investigate the binding profile of QSER1 in the chromatin, we performed ChIP-seq analysis (**Figure 4 and Figure EV4A)** using a validated QSER1 antibody (**Figures EV3G** and **Figure 6B**). We confirmed that QSER1 binds extensively on the chromatin of hESCs, as we detected an average of 12,461 peaks, near 7,942 genes. Importantly, a strong correlation was found between our datasets and Flag-tagged QSER1 ChIP-seq datasets previously published^40^ (**Figure EV4A**), although our ChIPseq retrieved less peaks, probably due to differences in the antibodies used (QSER1 versus Flag). The genomic distribution of our QSER1 ChIP-seq datasets revealed that around ∼65% of our QSER1 peaks lie on promoter regions, while the rest are spread among gene bodies and intergenic regions (**Figure EV4B**). Supporting its proposed role in safeguarding DNA methylation^40^, ∼48% of QSER1 peaks are located at CpG islands. However, the remaining peaks are found outside of CpG-rich regions, suggesting that QSER1 may have additional functions (**Figure EV4C**). To further characterize the chromatin context of QSER1 binding, we performed an overlap analysis using histone modification datasets from H1 hESCs (**Figure EV4D- E**). QSER1 peaks showed strong enrichment at regions marked by H3K27ac, a signature of active enhancers^81^, while a smaller subset overlapped with the repressive mark H3K27me3^81^ (**Figure EV4D–F**). The co-occupancy with distinct histone marks suggests the existence of functionally diverse QSER1-bound chromatin pools.

**Figure. 4.**
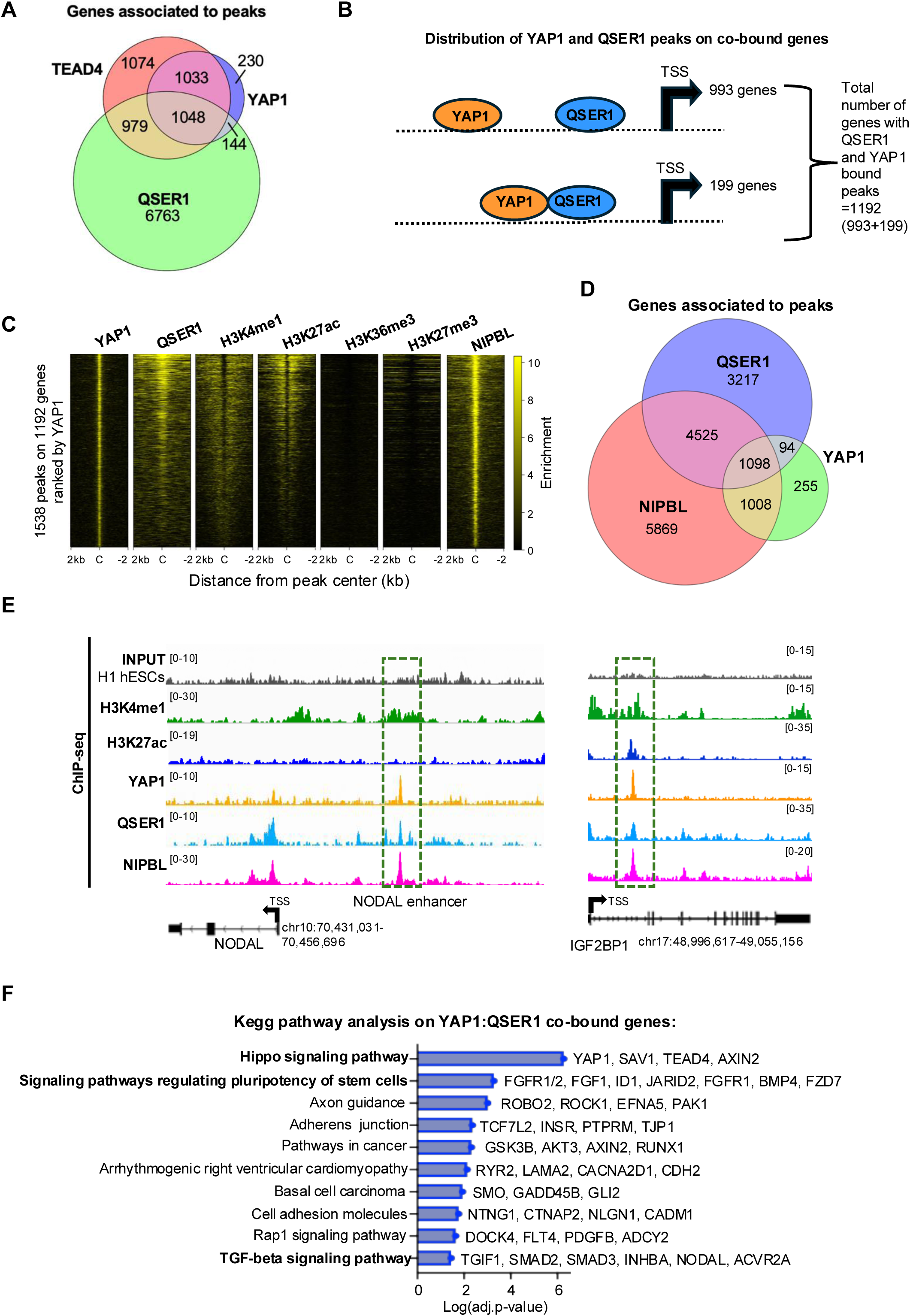
QSER1 binds YAP1: TEAD4 enhancers near developmental genes (A) Venn diagram shows the number of genes bound by QSER1, YAP1, and TEAD4 in hESCs (from ChIP-seq datasets). (B) Scheme shows YAP1 and QSER1 ChIP-seq peak distribution on co-bound genes in hESCs. Overlapping ovals denotate perfectly aligned peaks. Ovals separated by a dotted line represent distant annotated to the same gene. (C) Analysis of chromatin features associated to QSER1:YAP peaks. Heatmaps depict the correlation of QSER1:YAP co-bound sites with indicated histone modifications (source; ENCODE, Bernstein datasets). Peaks are organized from high to low YAP1 signal. (C = center of the peak, +/- 2Kb) . (D) Venn diagram shows the number of genes bound by QSER1, NIPBL (cohesin complex), and YAP1 (from ChIP-seq datasets) in hESCs. (E) IGV genome browser captures show YAP1 and QSER1 binding distribution at the indicated genes. H3K27ac and H3K4me1 histone marks are also shown. The coordinates of each genomic region are shown below each gene. TSS; transcriptional start site. The black arrow indicates the transcription direction. (F) Graph shows all enriched terms retrieved from Kegg Pathway analysis of the 1192 QSER1:YAP co-bound genes. Examples of genes included in each term are also indicated.

To gain insight into QSER1 potential activities, we used the professional services of a Molecular Modeling facility to analyze QSER1’s predicted structure through AlphaFold tools^79^. The results revealed that QSER1 is largely intrinsically disordered, with only two folded domains in the C-terminus: one of unknown function and a “bivalent Mical/EHBP Rab binding” (bMERB) domain^80^, a mostly helical domain, which is likely to mediate protein–protein interactions, including those with Rab GTPases (**Figure EV4G-H**). None of these domains are predicted to have enzymatic functions or catalytic activities. Notably, we found no evidence of a canonical DNA-binding domain, suggesting that QSER1 is likely recruited to chromatin through interactions with other DNA-bound proteins.

Given QSER1’s extensive intrinsically disordered regions and lack of predicted catalytic or DNA-binding domains, we propose that QSER1 functions as a scaffold or co-regulator that facilitates protein–protein interactions within chromatin-associated complexes.

### QSER1 co-localizes with YAP1:TEAD4 at developmental enhancers in hESCs

Next, we intersected the QSER1 ChIP-seq with YAP1 and TEAD4 ChIP-seq datasets we previously reported^37^. Results revealed substantial co-occupancy with 48.5% of YAP1-bound genes (1,192 out of 2,455) and 49% of TEAD4-bound genes (2,027 out of 4,134) also displaying QSER1 binding **(Figure 4A** and **Table EV3**). Nonetheless, QSER1 and YAP1 typically occupied distinct binding sites annotated to the same gene locus (**Figure 4B**). This is consistent with the general chromatin distribution of both proteins, with QSER1 peaks typically binding closer to promoters and YAP1 peaks closer to enhancers (**Figure EV4B**). Accordingly, most of the YAP peaks on the YAP1:QSER1 co-bound genes were enriched in typical enhancer marks H3K4me1 and H3K27ac^81^, and were co-occupied by cohesin, as shown by the profile of the NIPBL cohesin subunit^82^(**Figure 4C-4E**), suggesting that QSER1 may be brought into spatial proximity with YAP1 through chromatin looping. As stated above, genome-wide, ∼50% of QSER1 peaks overlapped CpG islands. In contrast, only 12% of QSER1:YAP1 co-bound peaks overlapped CpG islands (**Figure EV4C**). Together, these findings suggest that 1) QSER1’s role at YAP:TEAD-bound enhancers is likely independent of DNA methylation, and 2) that QSER1 recruitment to these sites may be mediated by TEAD or YAP1 tethering.

A KEGG pathway analysis^83^ of the 1192 co-bound genes revealed that the top significant enriched terms were “Signaling pathways in pluripotency” and “Hippo signaling pathway” (**Figure 4F**). Other significant categories include “Pathway in cancer”, which contain multiple Wnt and pluripotency genes, and “TGF-β signaling pathway” (**Figure 4F**). In these categories we found, in addition to *NODAL*, multiple TGF-β signaling genes like *INHBA*, which encoding homodimers form the Activin A ligand^84^ and its associated Activin receptor type IIA (*ACVR2A*), which both have important roles in early development^85^. We also found *TGIF1*, a critical inhibitor of the pathway^86^, and *SMAD2* and *SMAD3*, which are the NODAL/ACTIVIN/ TGF-β signaling effectors^87^. Among WNT-target genes we identified critical regulators of WNT/β-CATENIN signaling, including the gene codifying for the *GSK3b* kinase^88^, and *AXIN2*, a typical β-CATENIN target gene involved in restricting WNT activity^44^. We also found several members of the Frizzled family receptors, including *FZD7*, that mediate classical and non-classical WNT signaling^89^, and N-CADHERIN (codified by *CDH2* gene), a key molecule involved in epithelial-to-mesenchymal transitions, including those that occur during epiblast to PS transition in the embryo^90^. Moreover, critical pluripotency and germ-layer (i.e. *CDX2*, *SOX2*, *OTX2, KDM2B, PRDM14*), Hippo pathway (i.e. *TEAD4*, *AMOTL2*, *WWC1*), and Fibroblast Growth Factor- related genes (i.e. *FGFR1/2*, *FGFBP3*, *FGF1*) were also targeted by both proteins (**Table EV4**). Other categories, such as “Axon guidance”, “arrhythmogenic right ventricular cardiomyopathy” or “basal cell carcinoma” were also enriched, which likely reflect co-bound genes with broader or other context-specific functions. Taking together, we conclude that QSER1 and YAP1 co-occupy a common genetic program in hESCs enriched in pluripotency and lineage genes along with specific TGF- β/NODAL and WNT signaling genes.

### YAP1 recruits QSER1 to co-occupied developmental enhancers

YAP1 and QSER1 bind to 1192 genes in hESCs. From these, YAP1 and QSER1 peaks perfectly overlap on the same regulatory regions of 199 genes, while an additional 993 genes display YAP1 and QSER1 peaks at non-overlapping regulatory regions (**Figure 4B**). Here, we investigated the functional relationship between YAP1 and QSER1 on these genes. To do so, we analyzed the following three groups: (1) YAP1 and QSER1 co-occupied sites, such as those near *NODAL*, *CDX2* and *OTX2* genes; (2) YAP1 and QSER1 distal binding sites, like *SMAD2*, *SMARCA2*, and *INHBA* ; and (3) no YAP binding, which are genes bound by QSER1 that do not have YAP1 binding sites associated to the same gene, as seen in *PDX1* and *PAX6* (**Figure 5A-D and EV5A- B**). Then, we assessed QSER1 binding dynamics across these three types of regions in the presence and absence of YAP1, using ChIP-qPCR approaches. In YAP1 KO hESCs, we observed a significant decrease in QSER1 binding at regions co-occupied by YAP1, including *NODAL*, *CDX2*, *OTX2*, *SOX13*, and *SHB* genes, compared to WT (**Figure 5B**). In contrast, QSER1 binding at non-YAP1 target genes remained unaffected (**Figure EV5B)** . Interestingly, YAP1 deletion also reduced QSER1 levels on *SMAD2*, along with other distal QSER1-bound sites analyzed (**Figure 5D**). This further suggest a crosstalk between distal YAP1 and QSER1 bound regions, likely facilitated by cohesin-mediated distal interactions (**Figure 4C** and **4D**). On the other hand, QSER1 depletion (**Figure EV3F**) did not affect YAP1 binding on the analyzed regions (**Figure EV5C**). Altogether, these data show that YAP1 controls QSER1 recruitment on co-bound genes.

**Figure 5.**
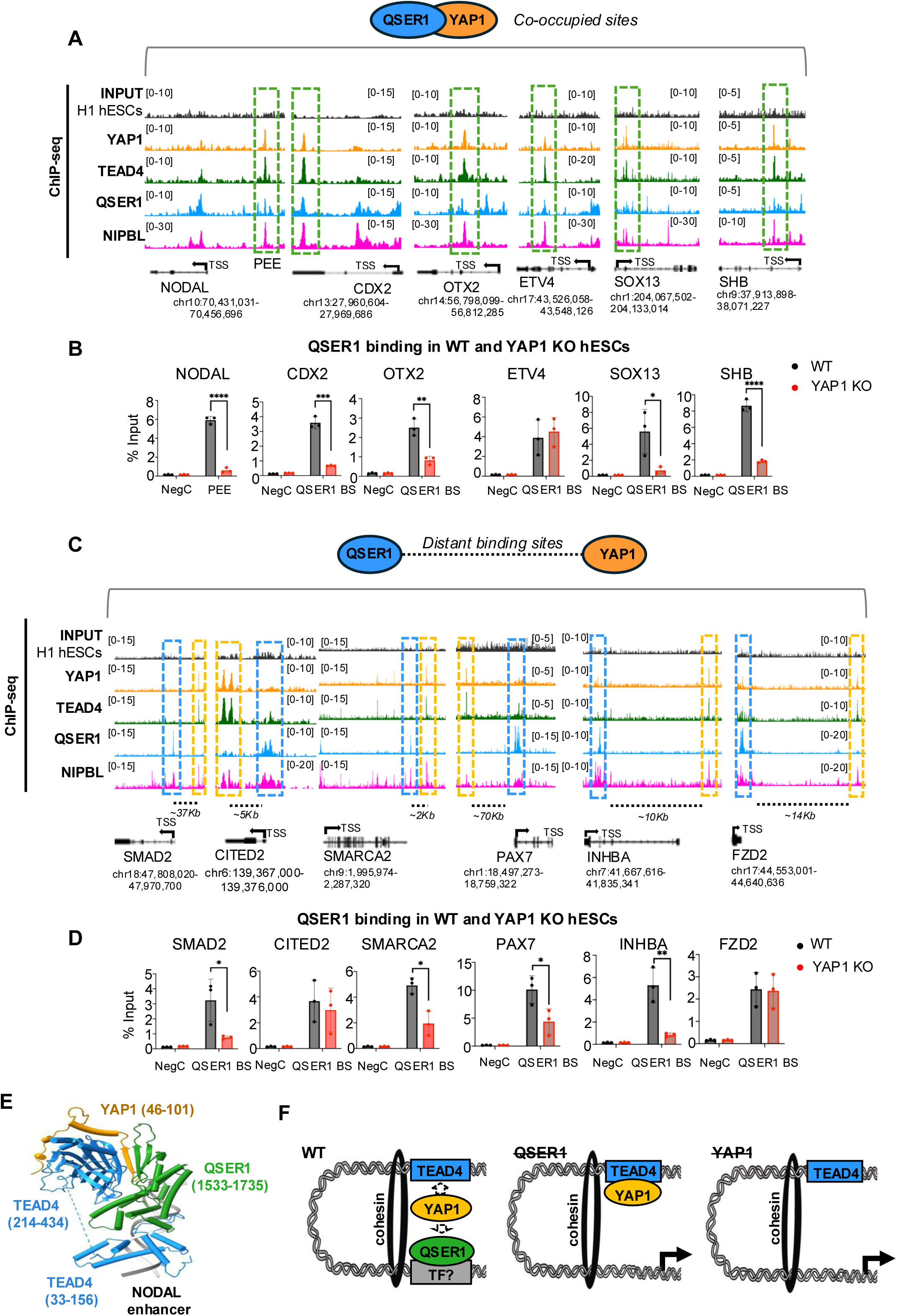
YAP1 tethers QSER1 to TEAD4 enhancers. (A) IGV genome browser captures show peak distribution of QSER1,YAP1, TEAD4, and NIPBL on indicated genes. Dotted green boxes enclose the co-occupied regions of QSER1 and YAP1. The coordinates of each genomic region are shown below each gene. TSS; transcriptional start site. The black arrow indicates the transcription direction. (B) Graphs show ChIP-qPCR analysis of QSER1 protein on the indicated genomic regions in WT and YAP1 KO hESCs. QSER1 BS: QSER1 binding site, depicted in **A**. NegC: Negative control region. (n=3) Data presented as mean ± SEM. Statistical analysis: Student’s t-test, ****p<0.0004 and ***p<0.001, *p<0.05 (C) IGV genome browser views showing the distal binding profile of YAP1, TEAD4, QSER1, and NIPBL on indicated genes. Blue and yellow boxes enclose QSER1 and YAP1 bound regions, respectively. Genomic coordinates corresponding to each locus are indicated below the gene tracks. TSS; transcriptional start site. Black arrow indicates transcription direction. (D) Graphs show ChIP-qPCR analysis of QSER1 protein on the distal binding sites in WT and YAP1 KO hESCs. QSER1 BS: QSER1 binding site, depicted in A. NegC: Negative control region. (n=3) Data presented as mean ± SEM. Statistical analysis: Student’s t-test, ****p<0.0004 and ***p<0.001, **p<0.01, *p<0.05 (E) AlphaFold3 Modeling of a complex of TEAD4-YAP1-QSER1 proteins with a cognate TEAD4 binding site on the NODAL enhancer. The domain labeling shows the boundaries of the interacting protein fragments in parentheses. Helices are shown as tubes and ß-strands as arrows for simplicity, and the flexible linker between the two TEAD4 domains is replaced by a dotted line. The two DNA strands are shown as gray ribbons and the main TEAD4 DNA binding helix is bound in the major groove of the cognate site. (F) Scheme summarizing interpretation of results. QSER1 is brought in proximity to TEAD-bound enhancers through YAP1. It is likely that the presence of cohesin facilitates these distal interactions. Since QSER1 does not have a DNA binding domain it is possible that binds DNA through other transcription factors (represented by the grey box; TF?).

**Figure 6.**
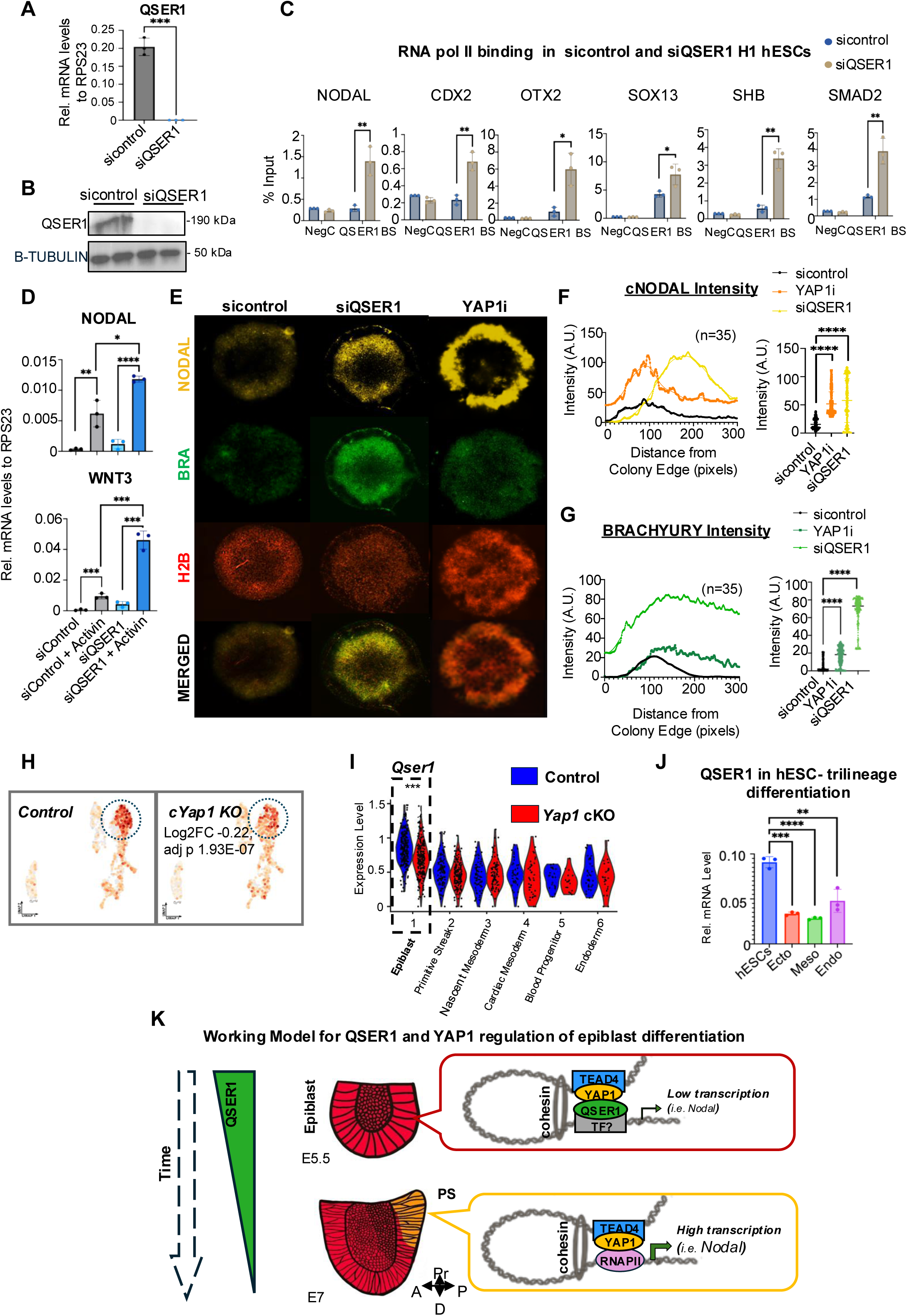
QSER1 regulates developmental signaling pathways in hESC-derived 2D gastruloids. (A) Graph shows QSER1 mRNA levels in hESCs transfected with siRNA control and siRNA against QSER1 for 72h. (n=3) Data presented as mean ± SEM. Statistical analysis: Student’s t-test, ***p<0.001 (B) Western blot of QSER1 protein levels, same conditions as **A**. (C) Graphs show ChIP-qPCR analysis of RNA polymerase II protein on the indicated genomic regions in sicontrol and siQSER1 hESCs. QSER1 BS: QSER1 binding site NegC: Negative control region. (n=3) Data presented as mean ± SEM. Statistical analysis: Student’s t-test, **p<0.01 and *p<0.05 (D) H1 hESCs were transfected with control or QSER1 siRNAs and left untreated or treated with Activin (=mesoderm inductor) for 24h. Graphs show RT-qPCR analysis of NODAL and WNT3 genes. (n=3) Data presented as mean ± SEM. Statistical analysis: Student’s t-test, ****p<0.0004, ***p<0.001, **p<0.01, *p<0.05 (E) Representative images of hESC-derived 2D-gastruloids with the genotypes indicated on the top are shown. To label the spatial expression of NODAL protein, an engineered line containing dual NODAL-citrine and H2B-RFP reporters was used (from Liu et al., 2022). A YAP1 inhibitor (0.5µM Dasatinib) was used to disrupt YAP1 activity (YAP1i). (F) Graphs show two types of quantification of citrine-Nodal levels (cNODAL). The plot on the left shows the intensity from the colony edge to the center of the micropattern, and the right plot shows the average intensity for cNODAL expression levels. (n=35) Data presented as mean ± SEM. Statistical analysis: Student’s t-test, ****p<0.0004 (G) Same as **F**, but quantification of immunostaining signal of Brachyury (BRA), a mesodermal fate marker. (H) UMAP from scRNAseq datasets of E7 embryos showing *Qser1* mRNA expression in control and *cYap1* KO embryos. Dotted circles highlight the epiblast cluster (see Figure 1). Differential *Qser1* expression in the epiblast of *Yap1* cKO versus control embryos is indicated. (I) Violin plot of *Qser1* from scRNAseq of E7 embryos showing expression levels in all clusters (Adjusted p-value is based on Bonferroni correction, ^∗^*p* < 0.05, ^∗∗^*p* < 0.001, ^∗∗∗^*p* < 0.0001.) (J) WT and YAP1 KO H1 hESCs were differentiated toward ectoderm (ecto), mesoderm (meso), or endoderm (endo) fates followed by RNAseq analysis (Stronati et al., 2022). Graph shows the expression of QSER1 from these datasets in the indicated conditions. (n=3) Data presented as mean ± SEM. Statistical analysis: Student’s t-test, ****p<0.0004 and ***p<0.001, **p<0.01). (K) Cooperative mechanism of YAP1 and QSER1 modulating gene expression of signaling genes in the mammalian epiblast. Two developmental stages are shown. QSER1 expression decreases as the epiblast transitions to PS, which allows RNAPII recruitment and increased transcription of genes, including *Nodal*. PS: primitive streak. Pr: proximal, A: anterior, P: posterior, D: Distal.

### Molecular Modeling Suggests YAP1 Acts as a Bridge Between TEAD4 and QSER1

To test whether YAP1 binding to TEAD proteins is required for QSER1 recruitment, we treated human pluripotent stem cells with GNE-7883^91^, a pan-TEAD inhibitor that allosterically disrupts the interaction between YAP1/TAZ and all TEAD paralogs by binding to the conserved TEAD lipid pocket. As observed with YAP1 depletion, treatment with GNE-7883 led to decreased CTGF and increased *NODAL* expression, indicating effective YAP1:TEAD complex disruption (**Figure EV5D).**

To investigate whether inhibition of YAP1 binding to TEAD affects QSER1 chromatin binding, we performed ChIP-qPCR at the NODAL enhancer, a region where QSER1, YAP1, and TEAD4 have been detected. GNE-7883 treatment significantly reduced both YAP1 and QSER1 occupancy at this site (**Figure EV5E**), supporting a model in which YAP1:TEAD complexes facilitate the recruitment of QSER1 to target enhancers.

To further explore the structural basis of this interaction, we used molecular modeling to predict the configuration of the YAP1:TEAD:QSER1 complex on the NODAL enhancer, using the DNA sequence encompassing the TEAD binding motif (see **Methods** for details). The resulting AlphaFold 3 modeling yielded a strong ipTM score of 0.68, consistent with a high-confidence trimeric complex on the DNA (**Figure 5E**). These structural predictions suggest that YAP1 functions as a molecular bridge: its residues 46–101 wrap around the second TEAD4 domain, and residues 50–60 are positioned in the interface between the bMERB domain of QSER1 and the YAP1 binding domain of TEAD4, acting like a bridge between QSER1 and TEAD4 (**Figure 5E and EV5F**).

Because of weaknesses in ipTM scoring^92^ particularly found in protein targets possessing large regions of intrinsically disordered residues (i.e QSER1 N-terminal residues 1 to 1320), we trimmed the interacting domains of each of the proteins to optimize the ipTM score (see boundaries enumerated in Methods). The resulting ipTM scoring for the TEAD4-YAP1-QSER1 complex showed a mean ipTM score of 0.61 for 25 models (standard deviation of 0.126) and strong conformational uniformity with a YAP1 β-strand bridging an exposed β-strand of TEAD4 and a β-strand of QSER1 (**Figure EV5F**). In contrast in the absence of YAP1, the TEAD4-QSER1 complex only yielded weak mean ipTM score of 0.31 (standard deviation of 0.27) for 25 models, which generated much more conformational heterogeneity in the location of QSER1 binding, and the resulting interfaces were weaker in general. This aligns with our ChIP-qPCR data, which show that YAP1 depletion leads to a marked reduction in QSER1 occupancy, while QSER1 loss does not affect YAP1 binding, which remains anchored via TEAD4 (**Figure 5B-5D and Figure EV5C**). Taken together, these structural predictions suggest that YAP1 functions as a molecular bridge, and in the absence of YAP1, QSER1 binding to YAP1:TEAD sites is compromised due to the loss of a stabilizing interface (**Figure 5F**).

### QSER1 depletion increase RNA polymerase II occupancy on YAP1-co-bound enhancers

In mouse embryos, a regulatory region known as the Proximal Epiblast Enhancer (PEE) is critical for NODAL expression in the PS during gastrulation^93^. Our previous studies identified that the PEE region is conserved in the human genome and that YAP1 represses this enhancer in hESCs^25^. In this study, we identified that QSER1 co-occupies this key enhancer of *NODAL*, located ∼13 kb upstream of the TSS in human cells. Thus, we speculate that QSER1 acts as a transcriptional co- repressor of YAP1 on this enhancer, to prevent excessive NODAL activation and subsequent premature differentiation of hESCs toward mesoendodermal lineages. To assess this, we analyzed RNA Polymerase II (RNAPII) binding to the *NODAL* enhancer. Our ChIP-qPCR analysis revealed that QSER1 depletion is sufficient to increase RNAPII abundance on the enhancer of *NODAL* (**Figure 6A-C and Figure EV6A**). Furthermore, we extend these analyses to the regions analyzed in **Figure 5B**. Our ChIP-qPCR analysis revealed a similar trend across most regions examined: QSER1-bound sites displayed increased RNAPII occupancy upon QSER1 depletion (**Figure 6C and EV6B**), suggesting that QSER1 may act to restrain transcriptional activity at the YAP1 co-occupied enhancers.

### QSER1 depletion induces NODAL signaling in 2D micropatterned gastruloids

Consistent with increased RNAPII occupancy at the *NODAL* enhancer, QSER1 depletion led to elevated *NODAL* transcript levels following mesoderm-inducing signals in hESCs (**Figure 6D** and see methods for details) and a similar transcriptional upregulation was observed for *WNT3* in response to QSER1 depletion (**Figure 6D**).

To gain more insight into the role of QSER1 in gastrulation, we used hESC-induced 2D- micropatterned gastruloids. In this system, hESCs are confined to circular micropatterns and exposed to BMP4 cytokine for 48 hours, which induces self-organizing embryonic germ layers (endoderm, mesoderm, and ectoderm) along the radial axis of the colony^94^. This organization arises from self-regulated signaling, with an ACTIVIN/NODAL gradient patterning mesendodermal fates in the outer ring (**Figure 6E**). In WT hESC patterns, similar to the PS in vivo, the mesoderm layer is identified by T/Brachyury expression, with its domain confined to the NODAL expression region^94,95^. To assess the function of QSER1, we transiently depleted QSER1 and analyzed a knock in Nodal-citrine:H2B-RFP hESC line that expresses the mature NODAL protein fused to a citrine tag^95^ (**Figure EV6C-D**). Consistent with the increase in the activity of the *NODAL* enhancer and transcript accumulation, QSER1-depleted gastruloids showed both elevated NODAL protein levels and an expanded expression domain compared to controls (**Figure 6E–F and Figure EV6D**). This was accompanied by a marked expansion of the mesodermal layer, as indicated by spreading of T/BRA expression toward the center of the colony (**Figure 6E, 6G and Figure EV6D**). Similar to what we previously shown for YAP1 KO phenotypes^25,37^, transiently blocking YAP1 activities using the inhibitor Dasatinib^96^ (YAP1i, **Figure EV6E**), was sufficient to induce NODAL protein expansion, along with mesoderm layer spreading, when compared with controls (**Figure 6E-G)**.

Together, these phenotypes reinforce the conclusion that QSER1 and YAP1 regulate a common transcriptional program during early lineage specification, restricting gene induction in response to developmental signals.

### QSER1 expression is enriched in pluripotent cells

We examined the expression pattern of *Qser1* across the early cell types of the E7 embryo. Unlike *Yap1*, which is homogenously distributed across pluripotent and differentiated cell types (**Figure 1**), *Qser1* is predominantly expressed in the epiblast, with reduced expression in more differentiated cell clusters (**Figure 6H-I**). A similar trend is observed in hESCs, where a significantly reduction in the mRNA levels of *Qser1* occurred during differentiation toward ectoderm, mesoderm and endoderm cell-fates, compared to pluripotent hESCs (**Figure 6J**). Furthermore, we identified a mild, although significant, reduction of *Qser1* levels in the *Yap1* cKO embryos compared to controls, further supporting the relevance of QSER1 in maintaining the epiblast (**Figure 6H-I**). Overall, we conclude that QSER1 is a specific co-factor of YAP1 in pluripotent cells required for the fine-tune regulation of gastrulation signaling genes (**Figure 6K**).

## DISCUSSION

In this study, we investigated the role of YAP1 in the differentiation of the mouse epiblast as well as the functional relationship with a new interactor in pluripotent cells, QSER1. Overall, our findings fill a long-standing gap in the field on defining the role of YAP1 in the differentiation of pluripotent cells, in vivo, and provide new mechanistic insights into how YAP1 regulates developmental genes.

Collectively, our findings support a model in which YAP1 plays a role in safeguarding the pluripotent epiblast state by modulating the threshold of mesendoderm-inducing signals. Specifically, YAP1 appears to actively suppress premature activation of NODAL and WNT pathways, thereby preventing ectopic or excessive primitive streak formation. This regulatory function is conserved across species and model systems, as evidenced by the convergence of in vivo mouse embryonic phenotypes with human stem cell–based gastrulation models^25,37,97^.

The PS is expanded in YAP1-deficient epiblasts at E7. This observation, together with impaired neuroectoderm differentiation, revealed by the cell population analysis at E7.75^28^ and morphological phenotyping at E8.25— suggests that YAP1 functions as a gatekeeper of lineage differentiation in the epiblast. Notably, enhanced PS specification and loss of anterior identity mirrors phenotypes caused by ectopic activation of WNT or NODAL signaling in the early embryo. For instance, excessive WNT signaling—such as that caused by activating mutations in *Lrp6* (a WNT co-receptor) or *Ctnnb1* (encoding β-CATENIN), or by deletion of the WNT inhibitor *Dkk1*—has been shown to disrupt neuroectoderm development and result in head truncation phenotypes⁹³^24^. Similarly, NODAL overexpression—whether by disruption of antagonists like *Lefty1/2*, or transcriptional repressors like *Tgif1/2*—lead to a variety of gastrulation defects, from expanded PS formation to failure to properly induce proper anterior ectodermal fates^39,86,98–101^. Thus, we speculate that, by limiting NODAL and WNT activity, YAP1 likely plays a critical role in ensuring proper spatial restriction of signaling domains during early patterning. However, our study did not include functional rescue experiments to test whether inhibition of NODAL or WNT signaling is sufficient to reduce PS differentiation or increase neural fold formation in *Yap1* mutant embryos. Additionally, YAP1 also modulates retinoic acid signaling at later stages in mesoderm populations of the embryos^28^. This suggest that YAP1 may regulate developmental signaling pathways in a cell-type dependent manner, contributing to embryonic development through multiple mechanisms. Future studies will be needed to determine the extent to which these pathways contribute to YAP1’s associated phenotypes during gastrulation.

Along these lines, we also observed potential non-autonomous effects of YAP1 deletion in extraembryonic populations. Extraembryonic tissues, including the anterior visceral endoderm (AVE) and extraembryonic ectoderm (ExE), are essential organizers of early embryonic patterning. These compartments act as key signaling hubs that influence epiblast behavior by secreting morphogens that regulate symmetry breaking, PS induction, and anterior–posterior (AP) axis formation. For instance, the ExE is a major source of BMP4, which induces *Wnt3* expression in adjacent epiblast cells to initiate PS formation, while the AVE secretes NODAL antagonists such as *Lefty1* and *Cer1* to restrict PS expansion and protect anterior fates^12,13,102,103^. Although our genetic strategy using Sox2-Cre deletes *Yap1* specifically in the epiblast, we observed significant changes in the composition of extraembryonic lineages in *Yap1* cKO embryos. These alterations may reflect non-cell-autonomous effects of YAP1 loss in the epiblast. However, it is important to mention that our *Yap1*^fl/fl^ embryos (control and *cKO*) are likely to be heterozygous for *Yap1* in extraembryonic tissues due to constitutive paternal allele deletion, through Sox2-Cre:Yap1^flox/+^ male breeders (see methods for details). Nonetheless, these observations raise the intriguing possibility that YAP1 may influence gastrulation not only through cell-intrinsic regulation, but also by modulating the epiblast–extraembryonic signaling axis. Future studies using lineage-specific Cre drivers to target YAP1 in extraembryonic compartments could help disentangle these effects and clarify whether YAP1 contributes to early patterning by affecting the formation or activity of extraembryonic cell populations.

Using a proximity labeling assay combined with *in silico* modeling and functional validation, we identified QSER1 as a novel co-factor of YAP1 in the regulation of a subset of developmental genes. QSER1 was recently characterized as a partner of TET enzymes and has been implicated in protecting CpG islands from DNA demethylation^40^. Our findings extend the understanding of QSER1’s functions, beyond DNA methylation. Our data showed that QSER1 depletion leads to NODAL de-repression and associates with increased RNA Polymerase II occupancy at YAP1- bound enhancers, pointing to a QSER1 contribution maintaining low transcriptional activity on certain genomic regions. Our in-silico analysis of the predicted structure revealed that QSER1 is an intrinsically disordered protein that lacks DNA-binding and catalytic domains. Based on these features and our findings, we hypothesize that QSER1 may act by stabilizing recruitment of transcriptional regulators, to fine-tune transcriptional levels, in accordance with the roles described for other intrinsically disordered proteins^104,105^.

Although YAP1 and QSER1 do not always co-occupy the same regulatory regions, our ChIP-qPCR analysis reveals that YAP1 influences QSER1 binding at both co-bound and distal sites, suggesting long-range regulatory effects. We propose that these distal regulatory regions, while separated linearly by several kilobases, are brought into spatial proximity through cohesin-mediated chromatin looping, enabling functional interaction between YAP1-bound and QSER1- bound elements. This model is supported by our findings showing extensive overlap between NIPBL, a core component of the cohesin complex^82^, and YAP1 peaks, suggesting that is frequently associated with loop anchor points, as previously reported^106^. This also highlights a key technical limitation of conventional ChIP-seq, which relies on identifying DNA fragments in close proximity to the protein–DNA crosslinking site, and it often fails to detect secondary DNA anchor points mediated indirectly through protein–protein interactions. To resolve this, future studies using chromatin conformation capture techniques, such as Hi-C^107^ or HiChIP^108^, will be necessary to determine how many YAP1- and QSER1-bound enhancers are physically connected, and how this spatial organization contributes to transcriptional regulation during early lineage specification. Finally, using hESC-based 2D gastruloid models, we showed that QSER1 depletion mimics the effects of YAP1 loss, and leads to enhanced NODAL and WNT signaling activities . Together, our results open new avenues to explore the role of QSER1 in in vivo models of gastrulation, and to investigate its broader functional interplay with YAP1 in other developmental contexts and disease states.

## MATERIALS AND METHODS

### Reagents and Tools Table

(Submitted in a separate file)

### Methods and Protocols hESC lines and culture

WT and YAP1 KO H1 hESCs were previously described^37^. The Nodal reporter (mCitrine::Nodal) hESC line was kindly provided by Dr. Aryeh Warmflash’s laboratory and described elsewhere^95^. The doxycycline-inducible Yap1-Myc-Bir2 H1 cell line was generated using a PiggyBac transposon system. In this process, YAP1 CDS, fused to a biotin ligase and Myc tag, was cloned into KA0717 plasmid, and co-transfected along the transactivator pPBCAG-rtTAM2-IN and transposase pCAG-PBase plasmids. These backbone plasmids were a gift from Dr. Kenjiro^37^. Transfected cells were isolated through G418 selection. Resistant clones were isolated and screened to confirm lack of leaking and robust expression of the construct upon doxycycline treatment. The doxycycline-inducible Yap1-Myc-Bir2 clonal cells were treated with 25pg/ml of Doxycycline to overexpress YAP1 without degradation. To confirm YAP1 deletion and overexpression, sanger sequencing and WB analysis were performed. All cell lines were tested regularly for mycoplasma using published procedures and primers^109^ and were maintained at 37°C and 5% CO2 in a humidity-controlled environment. All hESC lines were cultured in mTESR1 medium on Matrigel-coated tissue culture plates. The media was replaced every day. Colonies were split when they reached ∼70% confluence at a 1:20 ratio using 0.5M PBS/EDTA. For single cell dissociation experiments, all colonies were disaggregated using Accutase, cells were counted and seeded in the presence of Rock inhibitor for 24h. For directed differentiation to mesoderm, cells were treated with 50ng/ml Activin for 48h in mTESR1.

### Mice strains

All animal work complied with ethical regulations for animal testing and research and was conducted in accordance with IACUC approval by Temple University and followed all AAALAC guidelines (IACUC protocol #4973). All mice were obtained from Jackson Laboratory, including Yap1^fl/fl^ (*Yap1^tm1.1Dupa^*/J) and the Sox2-Cre mice (B6.Cg-Edil3<Tg(Sox2-cre)1Amc>/J). The generation of conditional knockout of YAP1 under Sox2-Cre were previously described^28^. In brief, *Yap1*^flox/flox^ mice were crossed with Sox2-Cre:Yap1^flox/+^ mice to generate conditional *Yap1* knockouts (Sox2-Cre:*Yap1*^fl/fl^) along with heterozygous littermate controls (*Yap1*^fl/fl^; Sox2-Cre-negative embryos). The *Yap1*^fl/fl^ embryos, even they are Sox2-Cre-negative, are likely to be heterozygous for *Yap1* in extraembryonic tissues due to paternal allele deletion, through Sox2- Cre:Yap1^flox/+^ males. This may have implications for interpreting potential effects on extraembryonic compartments (see discussion). Only males carrying the Sox2-Cre allele were used for breeding due to known maternal inheritance of Sox2-Cre^110^.

### Embryo isolation and Genotyping

We followed established protocol from our lab to isolate embryos and perform genotyping^28,30^ . Timed pregnancies were confirmed by the visualization of a vaginal plug at noon, which was considered E0.5. Embryos used in this study were isolated at E7-E9.5 for staining, qPCR, western blot and single-cell RNAseq analysis. The visceral yolk sac was collected for genotyping. Mice were genotyped using standard flox, cre, and SRY protocols (**Table EV5**).

### ChIP-qPCR and ChIP-seq experiments

ChIP experiments were conducted as previously described^21^ In brief, 1 x 10^6^ million cells were double crosslinked with 2mM di (N-succinimidyl) glutarate DSG (45min) and 1% formaldehyde (15 min) at RT. The cells were lysed and sonicated using a Qsonica Q700 prove sonicator (7 min ON, 1 min ON/OFF, 25% amp). Following sonication, the chromatin was centrifuged at max speed to remove the non-sonicated chromatin for 30 minutes. The chromatin extract incubated with the 1-5 ug primary antibody overnight at 4°C. After overnight incubation, protein A/G magnetic beads were added to the chromatin extracts for 4 hours at 4°C. The IPs were washed and reverse crosslinked with elution buffer overnight at 65°C. The next day, DNA purification was performed using the Qiaquick PCR purification kit. For ChIP-qPCR experiments, the amplification as performed on a Quant Studio 3 device using FAST SYBR green master mix or PowerUp SYBR green master mix for amplification. The DNA was eluted in 50-75µl and 2 µl was used for each reaction. For ChIP-seq experiments, the DNA was quantified using a Qubit Flex and the libraries were prepared using the NEBNext DNA Library kit. The quality of the libraries was analyzed using the Agilent 4150 Tapestation. Samples were sequenced using a P2 flow cell and NextSeq 1000 instrument. Primers for ChIP-qPCR and ChIP antibodies are listed in **Table EV6**.

### Biotinylation assay

The doxycycline (dox) -inducible Yap1-Myc-Bir2 H1 cell line was used to perform the proximity ligation assay. We performed this experiment using four different experimental groups (minus dox minus biotin, minus dox plus biotin, plus dox minus biotin, and plus dox plus biotion. To induce the protein construct, 25 pg/ml doxycycline was added to the media for 24 hours, followed by incubation with 50 mM biotin (or untreated) for 16 hours to label all proteins in proximity to YAP1. To identify nuclear interactors of YAP1, the nuclear fraction was isolated for analysis of biotin-labeled proteins. Briefly, nuclear lysis buffer (1 M Tris-HCl pH 7, 5 M NaCl, 1 M MgCl2, 10% NP-40, 1 M DTT, and 1x protease inhibitor) was added to the cells and incubated on ice for 10 minutes. The cells were then centrifuged at 500g for 5 minutes at 4°C. The supernatant was removed, and the pellet was washed with DPBS-/- and centrifuged again at 500g for 5 minutes at 4°C. The supernatant was discarded, and the pellet was snap-frozen and stored at -80°C before being shipped to the Proteomics Core at Sanford-Burnham-Prebys Medical Discovery Institute, where the pull-down and mass spectrometry were performed using an established protocol. Briefly, cells were lysed in 8M urea, 50 mM ammonium bicarbonate (ABC) and benzonase, then centri at 14,000 x g for 15 minutes. Disulfide bridges were reduced with 5 mM tris(2- carboxyethyl)phosphine (TCEP) at 30°C for 1 hour and alkylated with 15 mM iodoacetamide (IAA) in the dark at room temperature for 30 minutes at room temperature. Streptavidin-based affinity purification was performed on the Bravo AssayMap platform (Agilent) using AssayMap streptavidin cartridges (Agilent). Biotin-enriched peptides were reconstituted with 2% ACN, 0.1% FA, quantified by NanoDrop, and analyzed by LC-MS/MS using a Proxeon EASY-nanoLC system (ThermoFisher) coupled to a Orbitrap Fusion Lumos Tribid mass spectrometer (Thermo Fisher Scientific). Spectra were processed with MaxQuant software version 1.6.11.0 and searched against the Homo Sapiens Uniprot protein sequence database (downloaded in Jan 2020) and GPM cRAP sequences (commonly known protein contaminants). Limma software analysis was applied to identified differential enriched proteins in the four experimental conditions^111^. An initial list of 311 hits was obtained from proteins significantly enriched (FC>2, FDR<10%, with MS values in at least 2 replicates), in samples expressing YAP1-MYC-BIR2 construct treated with biotin (+dox,+biotin) versus those without biotin (+dox, -biotin). To refine the list of candidate interactors, the 311 initial hits were filtered based on fold change (Log2FC > 1.5) to exclude proteins that remained biotinylated in control conditions lacking the YAP1-MYC-BIR2 construct (−Dox, +Biotin) or both YAP1-MYC-BIR2 and biotin (−Dox, −Biotin). This filtering yielded a final subset of 83 proteins, which were then subjected to in silico protein–protein docking analysis to evaluate potential interactions with YAP1.

### Protein docking

Computation protein docking screening was performed on the list of 83 potential interactors from BioID screening of YAP1 and three positive controls (TEAD3, AMOT and LATS1). All protein structures were downloaded as PDB files from the AlphaFold human proteome dataset, which were shown to have a quality near to that of experimental methods as shown in the Critical Assessment of Techniques for Protein Structure Prediction (CASP14), making them a reliable option for in-silico protein studies^112–114^. HDock software was used to dock each protein to YAP1^115–119^. This software was chosen due to it being available as a standalone, downloadable package while also being one of the more accurate algorithms for protein docking according to the Critical Assessment of Predicted Interactions (CAPRI)^120^. HDOCK performs template-free, fast Fourier transform protein docking, which performs protein dockings *ab initio* without a homologous template. Since many of these complexes are novel interactions, template-free docking reduces possible bias from homologous templates. Fast Fourier transform protein docking converts the cartesian coordinates of each residue in both proteins into a three-dimensional grid with their associated values based on whether it is located on the surface, inside, or core of the protein^121^. One protein is kept static as the “receptor”, while the other protein is rotated around it as the “ligand”. HDock uses a 15-degree angle interval and a 1.2 Å translational interval, undergoing a total of 4392 rotations where one binding mode is kept per rotation. These output binding poses are then clustered with an RMSD of 5 Å, and the top ten structures are output as the final structures, in order of their scores from the HDock scoring function.

As a means of evaluating the stability of each of the docked structures, energy minimization calculations were performed using Amber software to provide an approximate ranking of these interactions^122^. This software was used due to its efficiency and ability to carry out quality simulations, especially those involving proteins and explicit solvent^123^. To prepare the docked protein files for use with Amber, they were first run on the H++ server, which predicts the protonation states of the histidine residues as well as the overall charge of the protein at biological pH^124^. The prepared PDB file from H++ was loaded in Amber with the ff19SB force field^125^. Ions were added to neutralize the system, then the protein was solvated using the OPC water model, which uses explicit solvation to model the aqueous solvent^126^. Ions were also randomly placed using the OPC model at a concentration of 150 mM to mimic the cellular environment^127^. Once the system was solvated and buffered, the energy minimization was run, which outputs a relaxed structure in its lowest energy conformation as well as the potential energy associated with this structure. This potential energy takes into account protein-protein interactions, protein-solvent interactions, and solvent-solvent interactions. While it does not give the binding free energy of the proteins, it outputs a potential energy value that can yield an approximate ranking and reflects the relative stability of the docked complexes. A more negative potential energy value from these calculations indicates that a complex is more stable, and the complexes with the lowest potential energy are more likely to represent biologically relevant interactions to be studied further.

### Treatment of inhibitors and Directed Differentiation of hESCs

For hESC experiments involving TEAD inhibition, cells were seeded and treated with 1µM GNE- 7883 for 24 hours before being collected for downstream analysis. For hESC experiments using YAP1 inhibitor DASATINIB, cells were seeded and treated with 0.5 µM for 72 hours. For directed differentiation to mesoderm, cells were treated with 50ng/ml Activin for 48h in mTESR1.

### Generation, Immunostaining, Imaging of 2D- hESC gastruloids

2D micropatterns were generated following a described protocol^94^. Briefly, micropattern chips were placed in a 6 well plate and coated with 5 µg/ml of laminin-521 in PBS +/+ for 2 hours at 37°C, then washed four times with PBS +/+. hESCs were dissociated with Accutase, centrifuged, and counted using the Invitrogen Countess 3. Cells were seeded with Rock inhibitor onto the glass chips. After 4 hours, the Rock inhibitor was removed and replaced with mTESR media for 12 hours. The following day, the cells were treated with 50 ng/ml of BMP4 for 48 hours.

For siRNA experiments, reverse transfections were performed directed onto the micropatterned glass chips for 24 hours. Total transfection length was 72 hours. Immunostaining of 2D gastruloids was performed following a stablished protocol^25^. Briefly, micropatterned chips were fixed with 4% formaldehyde for 20 minutes at room temperature (RT). The chips were then washed twice with PBS and permeabilized in a blocking solution (PBS with 0.1% Triton X-100, 3% Donkey Serum) for 45 minutes at room temperature. Primary antibodies were added to the blocking solution and incubated overnight at 4°C. The following day, the chips were washed twice with PBS and incubated with secondary antibodies and 1 mg/ml DAPI for 2 hours at room temperture. After incubation, the chips were washed three times with PBS and mounted on glass slides with mounting media. The micropatterns were imaged using an EVOS M7000 microscope with 10x and 20x objectives. Each channel was saved as a separate PNG file and analyzed with in-house MATLAB software, previously described^10^.

### Immunostaining, Imaging, and Quantification of hESCs

Glass chamber slides were coated with 5 µg/ml of laminin-521 in PBS +/+ for 2 hours at 37°C. Cells were dissociated using Accutase and seeded onto the coated slides. Once cells reached 60- 70% confluency, they were fixed with 4% formaldehyde for 15 minutes at room temperature.

Following fixation, cells were permeabilized with 0.5% Triton X-100 for 10 minutes and washed with PBS-/-. They were then incubated with blocking solution (0.5% PBS-Tween, 1%, BSA, 10% FBS) for 30 minutes at room temperature. Primary antibodies, diluted in blocking solution, were incubated overnight at 4°C. The following day, cells were washed and incubated with secondary antibodies and DAPI for 20 minutes at room temperature. Finally, cells were washed PBS and mounted with a glass coverslip.

Fluorescent intensity analysis was performed using the *ImageJ 1.53t* program. Images were taken using an EVOS M7000 microscope at 20 and 63x. A region of interest (ROI) was drawn using the freehand selection tool around the nucleus of a single cell using the DAPI channel. This ROI was then copied onto the RFP channel in the same exact position. Within the ROI, the sum of the intensity of the pixels was then measured in the DAPI and RFP channels individually. The RFP raw intensity was then normalized to the DAPI intensity of the same cell and multiplied by 100 to attain percent positivity. The fluorescence of 25 randomly selected individual cells were measured for each replicate. Pearson correlation coefficient was analyzed using the ZEN 3.0 (blue edition) software. Airyscan images of single cells were taken using the Zeiss LSM900 with Airyscan 2. DAPI, GFP, and RFP were imaged using the 405nm, 488nm, and 561nm lasers respectively. The ZEN 3.0 (blue edition) software was used to analyze the pearson correlation coefficient for the relationship between the colocalization of the GFP and RFP signal. In order to set the crosshairs for accurate analysis of positive correlation, the background colocalization of both the 488nm and 561nm lasers were determined by their colocalization with the unused 640nm laser for which no secondary antibody was utilized. Pearson correlation coefficient values above 0.7 are considered significant.

### RNA isolation, cDNA conversion, and qPCR

RNA extraction of hESCs was performed using the Quick RNA Extraction Zymo Kit. Mouse embryos were snap-frozen with Trizol in liquid nitrogen immediately after isolation. After genotyping, RNA extraction from mouse embryos was performed by pooling 3-4 embryos of each genotype. The Trizol procedure was applied following manufacture intstructions. In brief, following lysis with Trizol, chloroform and isopropanol were added to the samples for phase separation and RNA precipitation. Glycoblue was added to the RNA solution prior to resuspension in water. A total of 250-500 ng of RNA was reverse transcribed using the iScript Reverse Transcription Supermix. The cDNA samples were then amplified using FAST SYBR Green Master Mix or PowerUp SYBR Master Mix on a Quant Studio 3. All results were normalized to RPS23 and GAPDH unless otherwise stated, and the ΔΔCt method was used to calculate relative transcript abundance against the indicated references. Primers for cDNA amplification are listed in **Table EV6**.

### SiRNA transfections in hESCs

SiRNA transfections were performed using the Lipofectamine RNAiMAX transfection reagent following manufacturer’s instructions. Briefly, cells were dissociated using Accustase, and a mix of SiRNA, lipofectamine, and Opti-mem was added to the cells along with culture media and ROCK inhibitor. Transfection efficiency and downstream experiments were performed 72hr post-transfection.

### Protein Isolation, Co-immunoprecipitation (Co-IP) and Western Blots

For hESCs, cells were lysed using RIPA buffer containing protease and phosphatase inhibitors and left on ice for 20 minutes. For mouse embryos, after isolation, the embryos were snap-frozen in liquid nitrogen. Once the genotypes were confirmed, 8-10 embryos per genotype were pooled and lysed using RIPA buffer. Samples with RIPA were centrifuged at max speed for 10 minutes, and the supernatant was quantified using BCA assays. For Co-IP experiment, WT hESCs were washed with PBS-/- and lysed on ice for 20 minutes in IPH buffer (50mM Tris/HCl), 150 mM NaCl, 5mM EDTA, 0.5% Igepal). Lysates were then centrifuged at 13,000*g* for 10 minutes at 4 °C. The resulting supernatant was pre-cleared with agarose beads by rotating for 45 minutes at 4 °C. After separating the input, lysates were incubated overnight with primary antibodies at 4 °C with rotation. The following day, Protein A/G agarose beads were added and rotated for an additional 3 hours at 4 °C. Beads were washed three times with IPH buffer, and sample loading buffer was added. Samples were boiled at 95°C for 10 minutes and then loaded onto a gel. Following quantification, SDS-PAGE electrophoresis was performed with 10-40 µg of protein per sample for hESCs and 50-70 µg of protein per sample for mouse embryos. Wet transfer was carried out using 0.45 µm PVDF membranes. The membranes were blocked with 2% BSA in TBS-T for 1 hour at room temperature. Primary antibodies were diluted in 2% BSA in TBS-T and incubated overnight at 4°C. HRP-conjugated secondary antibodies and SuperSignal West Pico chemiluminescent substrate were used for protein detection on a BioRad Chemi-Doc Touch Detection System and LiCor Odyssey DLx Imager.

### Fast Protein Liquid Chromatography (FPLC)

Around 3 million cells of WT hESCs were lysed on ice for 30 minutes in Pierce^TM^ IP Lysis Buffer (Thermo Fisher) with 1× SIGMAFAST protease inhibitor cocktail. Lysates were centrifuged at 13,000*g* for 10 minutes at 4 °C and the protein concentration in the supernatant was determined using Pierce^TM^ 660nm Protein Assay Reagent (Thermo Fisher). 2 mg of protein were fractionated using an FPLC (ÄKTA pure FPLC, GE Healthcare), using a 1MDa Superose 6 Increase 10/300 GL column (#GE29-0915-96, Sigma-Aldrich) equilibrated in 1× PBS; 0.5-ml fractions were collected at a flow rate of 0.5 ml min^−1^. Then, fractions were concentrated to 75 μl with 3 kDa MWCO Amicon Ultra 0.5 centrifugal filter devices (#UFC500396, Merck Millipore) and used for immunoblotting. The molecular masses of the FPLC fractions were calibrated using gel filtration standards (#1511901, Bio-Rad Laboratories). Western blot intensities were analyzed using the ImageJ software. The intensity of the protein of interest was quantified and normalized to the intensity of the housekeeping protein.

### Single-cell RNAseq of E7 embryos

Procedures for single-cell analysis of mouse embryos during gastrulation followed protocols previously described by the Estaras lab^30,128^. For E7 analysis, multiple pregnancies were synchronized, and embryos were isolated at 7am seven days following the visualization of the vaginal plugs. Following genotyping, 3 cYap1 KO embryos (Sox2-Cre:Yapfl/fl) and 4 heterozygous controls (Sox2-Cre:Yapfl/+) were pooled and dissociated with Tryple for 5 minutes. The 10x Chromium Next GEM Single Cell 3’ GEM, Library & Gel Bead Kit v3.1 was used for library generation. Qubit and Agilent 4150 Tapestation were used for library quantification before pooling. Libraries were pooled and sequenced using NextSeq2000.

### Whole-mount immunostaining, Imaging, and Quantification of mouse embryos

After isolation, embryos were placed into a 48-well cell culture dish containing 4% formaldehyde and incubated for 45 minutes at room temperature. Following fixation, the embryos were washed three times with PBS and stored at 4°C. Once genotyping was confirmed, embryos of the same genotype were pooled into the same well. The embryos were permeabilized with 1% SDS in PBS for 15 minutes at room temperature while rocking. Afterward, the embryos were washed three times with PBS, and 0.5% Triton X-100 in PBS was added for 15 minutes at RT. The embryos were then washed three more times with PBS, and primary antibodies were added in blocking solution (PBS-T and 10% FBS). The following day, embryos were washed three times with PBS, and secondary antibodies, along with DAPI (1 mg/ml), were added for 2 hours at RT while rocking. The embryos were then mounted on glass slides with mounting media. Images were captured using the EVOS M7000. Immunofluorescence images were analyzed using the line tool in MATLAB to quantify the fluorescence intensity along the by proximal-to-distal axis of the embryo^25^. For the proximal-to-distal axis quantification; three “cup shaped” lines were drawn from the proximal posterior part of the epiblast to the most distal anterior part of the epiblast. The fluorescence signal of the three lines was averaged for each embryo. Each embryo image was rescaled to the same size, so distances are comparable. For the posterior-to-anterior quantification; the extraembryonic tissue and embryonic were located by eye. The embryo was divided in six bins by drawing five parallel straight lines from the boundary between the extraembryonic tissue down to the distal tip of the epiblast. The average fluorescence signal of each bin was plotted in the graph (from posterior to anterior). The average of the intensity profiles from 3 embryos per condition were plotted. Brightfield images were also taken after each embryo isolation, and phenotyping analysis was conducted. The perimeter of the epiblast was quantified using ImageJ, and the number of cells per embryo was calculated by trypsinizing the cells and counting them using a Countess 3 device.

### Modeling AlphaFold Proteins and Protein-DNA complexes

The AlphaFold3 (AF3) and AlphaFold2 (AF2) programs were utilized^79^. Protein sequences and domain boundaries were refined from Uniprot entries for the following proteins: TEAD4_HUMAN (Q15561), YAP1_HUMAN (P46937), and QSER1_HUMAN (Q2KHR3). AF3 and AF2 modeling predicted QSER1 structure and identified domains. The AF3 program was utilized to model transcription factors bound to DNA forming a variety of complexes. Given the experimental evidence for transcription factor binding to the Nodal enhancer presented in this work, a 36 base-pair portion of this DNA region was chosen as it contained a single TEAD4 cognate binding site (sequence T<u>GCATTCC</u>CCACTAACATCAAAAAGCCTGGGAGAGC,cognate binding site underlined) To make a model for this specific portion of the enhancer, a TEAD4 protein monomer, and a single peptide of YAP1, and various portions of the QSER1 protein were utilized. In all cases, the models produced had the DNA-binding domain of TEAD4 positioned precisely to allow extensive binding contacts between the main DNA-binding helix to the major groove of the cognate DNA sequence. In comparing TEAD4 binding to its cognate sequence in known structures, the AF3 model had 8 hydrogen bonds to DNA compared to 10 hydrogen bonds found in the known X-ray crystal structure PDB code 5GZB, with 6 amino acids in common to both structures.

TEAD4 has an N-terminal DNA binding domain followed by a flexible unstructured linker which connects to a second folded effector domain in the C-terminal half of the protein. The cognate DNA sequence strictly governed the position of the N-terminal TEAD4 DNA binding domain, while the flexible linker allowed many different positions for the C-terminal domains among the various models sampled. When YAP1 is present (especially residues 46-101) it wraps around the second TEAD4 domain and acts like “molecular glue” to bridge TEAD4 binding to the QSER1- bMERB domain found at the C-terminus. A beta strand from YAP1 inserts between the TEAD4 domain and the QSER1-bMERB domain to form a tighter interface that AF3 predicts to be more stable. While previous crystal structures have shown mouse (3JUA) and human (8A8R) YAP1 wrapping around a single TEAD domain, these AF3 models demonstrate a YAP1 peptide inserting between TEAD4 and QSER1. Aligning and super-positioning of protein-DNA complexes, analysis of interfaces and contacts, and figure preparation were done with ChimeraX^129^.

### Quantitative scoring and N-terminal trimming of proteins used in modeling

AF3 modeling was first performed with full-length canonical protein sequences from Uniprot. To optimize AF3 modeling and scores, subsequent models were built with protein sequences that trim away flexible unstructured N-terminal tails as follows: TEAD4(1-32), YAP1(1-45), and QSER1 (1-1532). The final domain boundaries are shown as residue numbers in the color-coded labels of the model **Figure 5E**. To quantitatively compare the scoring of protein complexes in the presence and absence of YAP1(46-101), the DNA was omitted and the ColabFold version^130^ of AlphaFold2 was used with higher sampling (n=25 models for each protein complex) and total ipTM scores were compared with and without the YAP1 peptide. The geometric mean of the ipTM scores and standard deviation (Excel STDEV.P function for the whole population) were calculated for each set of 25 models. Structural and conformational homogeneity was assessed by aligning the 25 models on TEAD4 domains, and visual inspection of the homogeneity of YAP1 and or QSER1 positioning.

### scRNAseq analysis of E7 embryos

Single cell sequencing reads were counted individually for each sample using cellranger (v7.1.0) with default parameter on mm10 mice genome. Detection of doublet and cells contaminated with ambient RNA were assessed using scrublet and soupX (v1.6.2), in Python and R, respectively. Low-quality cells, including doublets, cells with over 10% mitochondrial content, and those expressing fewer than 5000 RNA features, were filtered out. Most analysis were performed using the Seurat R package (v4.3.0), including quality control plots generated with FeatureScatter and VlnPlot Seurat’ functions. Samples were normalized and variance stabilized using the SCTransform function in Seurat, regressing out variables including nCount_RNA, percent.mt, percent.rb, S.Score, and G2M.Score. Principal component analysis (PCA) was run with 19 principal components for both samples. The samples were then integrated using Seurat’s SelectIntegrationFeatures, PrepSCTIntegration, FindIntegrationAnchors, and IntegrateData functions with 3000 selected features and the SCTransform normalization method. The populations were annotated based on the differential expression of known markers tested against all other clusters combined. Selected markers with significant adjusted p-values in the assigned population were used to generate the heatmap in **Figure 1B**. For example, Sox2 is significantly upregulated in the epiblast cluster relative to all other clusters. For visualization of gene expression in UMAP, and heatmap, the SCT, and RNA assay were used, respectively. Differential gene expression analysis for each cluster between genotypes was performed on the RNA assay using the FindMarkers Seurat function after log normalization and scaling. Genes were considered differentially expressed when adjusted p-value, based on Bonferroni correction was below 0.05. Cell type count differences between genotypes for each cell types were assessed by randomly downsampling cYap1 KO samples 100 times to the same number of cells as the WT sample (788 cells). For each iteration, the number of cells in each cluster was counted. Chi-squared tests with Bonferroni correction were used to assessed significant differences in cell counts between genotypes across clusters. Pathway analysis was performed using the SCPA R package (v1.5.4) with curated gene lists selected from the MSigDB C2 database.

### ChIP-seq analysis

ChIP fragments were sequenced in an Illumina NextSeq 1000 sequencer in SE configuration. Adapter and poor-quality sequences were trimmed, and cleaned, using fastp (v0.23.2). Reads were aligned to the Human hg38 genome assembly (GRCh38)using bowtie2 (v2.2.5) withparameters ‘--phred33 -q --no-unal’. Reads with a mapping quality score (MAPQ) below 20, unmapped reads, secondary alignments, and reads failing quality checks were removed using SAMtools (v1.16.1), and duplicated reads were removed using Picard’s MarkDuplicates function. Peaks were identified using getDifferentialPeaksReplicates.pl from HOMER (v4.11) with style ‘-factor’ parameter and input sample as background reference. Peaks overlapping were assessed using SRplot (***ref: PMID: 37943830***). Differential peak enrichment between genotypes was conducted using rgt-THOR (***ref: PMID: 27484474***) with parameters ‘--merge --deadzones’ to combine peaks, and exclude regions from the hg38 blacklist. In addition to differential peak calls, rgt-THOR generated a bigwig file for visualization. Peaks, and differential peaks, were assigned to genes using the nearest TSS method with ChIPseeker (v1.18.0). Two independent sets of histone modification ChIP-seq samples for H3K27me3, H3K4me1, H3K36me3, and H3K27ac were obtained from ENCODE, generated by the labs of Bing Ren (GSE16256) and Bradley Bernstein (GSE29611). Peak and bigwig coverage files were directly retrieved from ENCODE for downstream analysis. CpG islands were retrieved from the hg38 UCSC genome annotation database. Heatmaps of ChIP-seq signals were generated using deepTools’ computeMatrix and plotHeatmap functions. Gene ontology analysis was performed using the enrichR R package (v3.2) using the GO_Biological_Process_2023 library and visualized with ggplot2.

## DATA AVAILABILITY

ChIP-Seq and scRNAseq data were deposited in GEO under the accession no. GSE280805 and GSE280804, respectively.

## AUTHOR CONTRIBUTIONS

Conceptualization, C.E. and E.A.,; methodology, C.E. and E.A.; investigation, C.E., E.A., A.D., E.M., O.M.P, C.C-F., J.F.G, H.M.C., P.R.F., M.Z., S.K., J.W.E, and M.D.A ; formal analysis, J.E.H, M.D.K., T.R., and N.A; visualization, C.E., E.A., A.D., E.M., O.M.P, M.D.A, J.E.H, T.R., and N.A; writing – original draft, C.E. and E.A.; writing – review & editing, C.E. and E.A.; funding acquisition, C.E. and E.A.

## Supporting information

Tabl3 EV2

## ACKNOWLEDGMENTS

We acknowledge previous members of the Estaras lab for discussions and experimental support, including Dr. Eleonora Stronati and Arantza Larrinaga-Zamanillo. Also, thanks to Dr. Simon, Dr. Kishore, and Dr. Garikipati’s laboratories at Aging+Cardiovascular Discovery Center for discussions and feedback. Thanks to Dr. Kishore’s lab for their generosity sharing reagents. Thanks to Alex Morris (Fox Chase Cancer Center) for initial analysis on scRNAseq datasets and discussion. Dr. Jonathan Whetstine’s lab and the Genomics Facility at the Fox Chase Cancer Center for technical support for single-cell experiments. We acknowledge support from Fox Chase Molecular Modeling Facility, funded in part by the NIH Cancer Center Support Grant P30 CA006927. Thanks to Sanford Burnham Prebys Proteomics and Bioinformatics Core for help with the biotinylation assay and bioinformatics analysis, funded in part by the NIH Cancer Center Support P30 CA030199. This work was funded by the NIH/NICHD R01 HD106969 (to C.E), NIH/NINDS R01NS119699 (to N.A), NIH/NHLBI 5T32HL091804 (to E.A and E.M), NIH/NICHD F31HD113419 (to E.A), and NIH/NIGMS T32 GM142606 (to A.D).

Table EV1 – DEGs in E7 mouse embryos Yap1 cKO compared to control

Table EV2 – BioID hits from hESCs

Table EV3 – Venn diagram from ChIPseq for YAP1, QSER1, and TEAD4 in WT H1 hESCs

Table EV4 – Kegg pathway of QSER1:YAP1 peaks

Table EV5 – PCR primers for genotyping and PCR conditions

Table EV6 – qPCR primers

Expanded View Figure 1.

(A) Gel shows expected bands from flox genotyping using a Yap1Flox:cre system. Adapted from Abraham et al., 2025.

(B) Gels show the genotype of breeding partners. Note that only males carry the Sox2-cre allele to avoid maternal inheritance of Cre activity.

(C) Same-day genotyping for flox and Cre for fresh-embryo sequencing was performed from the yolk sacs of 14 embryos, simultaneously isolated from 2 pregnant dams. Four controls, indicated in red triangles, and three *Yap1* cKO embryos (floxflox/cre+), shown in blue circles, were pooled and processed for scRNAseq.

(D) Genotyping of SRY (sex identity) in the 14 embryos isolated for the experimental design of the scRNAseq experiment.

(E) Violin plot of *Yap1* and *Wwtr1* (TAZ) from scRNAseq expression levels in all clusters comparing *Yap1* cKO to control.

(F) Graphs show RT-qPCR analysis of *Yap1* and its target gene, Ccn2 (CTGF), in E7.5 *Yap1* cKO and control embryos. (n=10) Data are presented as mean ± SEM. Statistical analysis: Student’s t-test, **p<0.01 and ***p<0.001.

(G) Graphs display cell cycle S and G2M scores in control and *Yap1* cKO embryos from scRNAseq analysis.

(H) Bright-field images of control and *Yap1* cKO E7 embryos. Graphs show cell number quantification per embryo (left) and the size of the epiblast (right) in control and *Yap1* cKO embryos. (n=8-10 embryos) Data are presented as mean ± SEM. Statistical analysis:

Student’s t-test. Scale Bar 250µm.

(I) Single Cell Pathway analysis was applied to DEGs. Terms related to TGFb and Wnt signaling pathways significantly enriched (q-value>1.4, adj. p-value <0.05) in the epiblast are shown.

(J) Full western blot of nuclear extracts of E7 embryos shown in **Figure 1G**. C: control embryos and Y: *Yap1 cKO* embryos. Red Arrows indicate bands shown in main Figure; SMAD2/3 (mw: 55kDa), HISTONE H3 (mw: 15kDa), GAPDH (37kDa), Β-CATENIN

(mw: 90 kDa).

(K) Western blot of whole embryo lysates of E7 control and *Yap1* cKO embryos. Pooled embryos numbers are indicated above each lane, along with the makers analyzed and on the right is the full blots. Red Arrows indicate bands that were cropped; SMAD2/3 (mw: 55kDa), GAPDH (37kDa), and Β-CATENIN (mw: 90 kDa).

Expanded View Figure 2.

(A) Bar graph showing the percentage of cells assigned to each cluster of control and Yap1 cKO embryos. All populations are shown. Statistical analysis: Chi-test, * < 0.05, ** < 0.001,*** < 0.0001.

(B) Merged images of whole-mount immunostaining of BRACHYURY (T; green) and DAPI (blue) in E7.5 control and *Yap1* cKO embryos. The experiment was repeated three times with different litters with consistent results. See also **Figure 2B**. Scale Bar 250µm.

(C) Bright-field images of control and *Yap1* cKO at E8.25 and E9. hf: head fold, ng: neural groove, cc: cardiac crescent, nt: neural tube, hrt: heart, s: somites. Scale bar 250µm.

(D) Table showing the phenotyping description of control and *Yap1* cKO embryos at E8.25 (n=15).

(E) Percentage of cells from E7.75 control and *Yap1* cKO scRNA-seq datasets previously published by our lab (Abraham et al., Cell Reports 2025). Clusters marked with an asterisk (*) are significantly different. Statistical analysis: Chi-test, ***p < 0.0001.

(F) Western blot of whole embryo lysates of control and *Yap1* cKO at E7. Embryos pooled per lane are indicated, along with the markers. The uncropped blots are represented below. The red arrows highlight the molecular weight. C: control Y: *Yap1* cKO Eomes (mw: 70kDa) and Gapdh (mw: 37kDa) *Same western blot from **Figure EV1K.**

(G) qPCR of listed markers in E7.5 embryos. (n=10) Data are presented as mean ± SEM. Statistical analysis: Student’s t-test, ^∗∗^*p* < 0.001.

Expanded View Figure 3.

(A) Uncropped western blot from **Figure 3B**. Red arrowheads indicate bands shown in the main figure.

(B) Uncropped western blot from **Figure 3C**. The dotted square highlights the bands shown in the main figure.

(C) Co-immunoprecipitation (Co-IP) experiment was performed in the doxycycline-inducible YAP1-Myc-Bir2 clonal hESC line in the presence and absence of Doxycycline, as indicated. The YAP1 antibody was used for immunoprecipitation, and western blot analysis was performed using a QSER1, TEAD4, and YAP1 antibodies. 10% of total lysate was loaded as input. Full western blot shown below.

(D) ChIP-qPCR analysis was performed in the clonal YAP1-Myc-Bir2 hESC line, in the absence and presence of 25pg/ml of Doxycycline (Dox). ChIPs using c-myc and YAP1 antibodies were carried out. The genomic regions analyzed are indicated at the bottom. NegC, negative control region. (n=4) Data are presented as mean ± SEM. Statistical analysis: Student’s t-test, ^∗∗^*p* < 0.01 and ^∗∗∗^*p <* 0.001.

(E) Table depicting experimental conditions used for the BioID2 assay and replicates. Raw MS/MS counts (without normalization) are shown for the indicated proteins in each replicate. Dox: doxycycline.

(F) Immunostaining of endogenous YAP1 (green) and QSER1 (red) in WT hESCs. Representative 63x magnified images of single cells are shown. Scale bar 2µm. (n=25) On the right, Pearson correlation coefficient (r) was calculated for YAP1 and QSER1 signals in each cell. A threshold of r >0.7 (indicated by the red dashed line) shows a strong positive correlation.

(G) Immunostaining of QSER1 (red) in sicontrol and siQSER1 in H1 hESCs and quantification of cells positive for QSER1, relative to DAPI (blue). Scale bar 50µm. (n=25) Data are presented as mean ± SEM. Statistical analysis: Student’s t test, \*\*\*\**p*

*<* 0.0001.

(H) Immunostaining of YAP1 (green) in WT and YAP1 KO H1 hESCs and quantification of cells positive for YAP1, relative to DAPI (blue). Scale bar 10µm. (n=25) Data are presented as mean ± SEM. Statistical analysis: Student’s t test, \*\*\*\**p <* 0.0001.

(I) Size-exclusion chromatography was performed on nuclear extracts from WT H1 hESCs. The elution fractions analyzed are indicated on the top. The approximate MW range covered by these fractions is indicated at the bottom. Western blot of QSER1 and YAP1 was performed. Relative intensity was quantified. Complete blots are shown on the right. * shows a band that appeared in both the QSER1 and YAP1 channels and was discarded from quantifications.

Expanded View 4.

(A) Intersection analysis was carried out between our QSER1 ChIP-seq (using QSER1 antibody) and previously published Flag-tagged QSER1 ChIPseq in H1 hESCs (using

FLAG antibody) (Dixon *et al*., 2021). Heatmaps and Venn Diagram show the binding correlation and number of co-bound genes in the two datasets.

(B) Genomic distribution of QSER1, TEAD4, and YAP1 peaks in WT H1 hESCs from ChIP- seq datasets.

(C) Graph shows the percentage of QSER1 peaks (all peaks) and QSER1:YAP1 co-bound peaks (199) associated to CpG islands in hESCs.

(D) Heatmap shows correlation of QSER1 binding, indicated histone marks (source, ENCODE, Bernstein datasets), and EZH2. The heatmap is ranked by QSER1 peak signal and the number of peaks are shown (C = center of the peak, +/- 2kb).

(E) Table shows the number of overlapping QSER1:H3K4me1, QSER1:H3K27ac, QSER1:H3K36me3, and QSER1:H3K27me3 peaks, genome-wide, using two different histone ChIP-seq datasets (Ren and Bernstein).

(F) IGV genome browser captures show QSER1, H3K27me3, and H3K27ac at indicated genes TSS: transcription start site.

(G) QSER1 predicted structure based on AF modeling. Most of the protein comprises an intrinsically disordered region (IDR), especially the first three quarters (residues 1 to 1319, orange). Following the large IDR, there are two folded domains. The first is a domain of unknown function (light blue) that is mostly composed of ß-strands, and the second has been designated as a “bivalent Mical/EHBP Rab binding” or bMERB domain (magenta, residues 1535-1735) and is largely helical with three ß-strands in the middle. These two folded domains are separated by 90 residues of a flexible disordered linker (also orange). Below the domain arrangement is a scale bar to denote residue numbers and a bar that represents the AlphaFold2 confidence score (plDDT) with a color key below that shows that darker blue reflects higher confidence.

(H) Schematic representation of IDR and AlphaFold predictions of QSER1 folded domain structures. Each is color-coded to match the domains shown above. While the relative orientation of the two domains is the most frequently predicted conformation, the linker between them is highly flexible.

Expanded View 5.

(A) IGV genome browser snapshots show more examples of distribution of QSER1,YAP1, TEAD4, and NIPBL on indicated genes.

(B) Graphs show ChIP-qPCR analysis of QSER1 protein on the indicated genomic regions in WT and YAP1 KO hESCs. QSER1 BS: QSER1 binding site. NegC: Negative control region. (n=3) Data presented as mean ± SEM. Statistical analysis: Student’s t-test.

(C) Graphs show ChIP-qPCR analysis of YAP1 protein on the indicated genomic regions and conditions in sicontrol and siQSER1 conditions. NegC: Negative control region. (n=3) Data presented as mean ± SEM. Statistical analysis: Student’s t-test.

(D) RT-qPCR of gene expression of CTGF (downstream gene of the Hippo signaling pathway) and NODAL in WT H1 hESCs treated with or without 5µM GNE-7883 TEAD inhibitor (TEADi) (n=3) Data presented as mean ± SEM. Statistical analysis: Student’s t- test, **p<0.01.

(E) Graph of ChIP-qPCR of TEAD4, YAP1, and QSER1 at enhancer of the NODAL gene in untreated and TEADi treated cells. NegC: Negative control region. (n=3) Data presented as mean ± SEM. Statistical analysis: Student’s t-test, **p<0.01.

(F) Molecular modeling of TEAD4 (blue), YAP1 (orange), and QSER1 (green) using AlphaFold3 showing that YAP1 residues 50-60 are tightly bound to QSER1 residues

1613-1623 (7 hydrogen bonds) and TEAD4 residues 340-349 (5 hydrogen bonds, shown as dotted lines). Top ipTM scores for this complex are 0.68, reflecting a high confidence in the conformation of this model.

Expanded View 6.

(A) Full uncropped blot of **Figure 6B**. Blotted against QSER1 (mw: 190Kda) and beta- TUBLIN (mw: 50 Kda). Red arrow indicates the band that was cropped. Sic: sicontrol and SiQ: siQSER1.

(B) Graphs show ChIP-qPCR analysis of RNA polymerase II protein on the indicated genomic regions in sicontrol and siQSER1 hESCs. NegC: Negative control region and QSER1 BS: QSER1 binding site. (n=3) Data presented as mean ± SEM. Statistical analysis: Student’s t-test.

(C) Scheme of the Nodal-citrine: H2B-RFP hESC construct with representative fluorescent images of hESCs under basal conditions.

(D) Additional images of hESC 2D gastruloids shown in **Figure 6E**.

(E) Graphs show RT-qPCR analysis of YAP1-target genes CTGF and CYR61 in hESCs untreated and treated with the YAP1 inhibitor (YAPi) DASATINIB for 72h treatment. (n=3) Data presented as mean ± SEM. Statistical analysis: Student’s t-test, *p<0.05

## REFERENCES

1. Wu Z, Guan KL. Hippo Signaling in Embryogenesis and Development. Trends Biochem Sci. Jan 2021;46(1):51–63. doi:10.1016/j.tibs.2020.08.008

2. Hossain Z, Ali SM, Ko HL, et al. Glomerulocystic kidney disease in mice with a targeted inactivation of Wwtr1. Proc Natl Acad Sci U S A. Jan 30 2007;104(5):1631–6. doi:10.1073/pnas.0605266104

3. Morin-Kensicki EM, Boone BN, Howell M, et al. Defects in yolk sac vasculogenesis, chorioallantoic fusion, and embryonic axis elongation in mice with targeted disruption of Yap65. Mol Cell Biol. Jan 2006;26(1):77–87. doi:10.1128/MCB.26.1.77-87.2006

4. Zhong Z, Jiao Z, Yu FX. The Hippo signaling pathway in development and regeneration. Cell Rep. Mar 26 2024;43(3):113926. doi:10.1016/j.celrep.2024.113926

5. Dong J, Feldmann G, Huang J, et al. Elucidation of a universal size-control mechanism in Drosophila and mammals. Cell. Sep 21 2007;130(6):1120–33. doi:10.1016/j.cell.2007.07.019

6. Kowalczyk W, Romanelli L, Atkins M, et al. Hippo signaling instructs ectopic but not normal organ growth. Science. Nov 18 2022;378(6621):eabg3679. doi:10.1126/science.abg3679

7. Lu L, Finegold MJ, Johnson RL. Hippo pathway coactivators Yap and Taz are required to coordinate mammalian liver regeneration. Exp Mol Med. Jan 5 2018;50(1):e423. doi:10.1038/emm.2017.205

8. Yu C, Ji SY, Dang YJ, et al. Oocyte-expressed yes-associated protein is a key activator of the early zygotic genome in mouse. Cell Res. Mar 2016;26(3):275–87. doi:10.1038/cr.2016.20

9. Nishioka N, Inoue K, Adachi K, et al. The Hippo signaling pathway components Lats and Yap pattern Tead4 activity to distinguish mouse trophectoderm from inner cell mass. Dev Cell. Mar 2009;16(3):398–410. doi:10.1016/j.devcel.2009.02.003

10. Ghimire S, Mantziou V, Moris N, Arias AM. Human gastrulation: The embryo and its models. Dev Biol. Jan 2021;doi:10.1016/j.ydbio.2021.01.006

11. Wang R, Yang X, Chen J, et al. Time space and single-cell resolved tissue lineage trajectories and laterality of body plan at gastrulation. Nat Commun. Sep 14 2023;14(1):5675. doi:10.1038/s41467-023-41482-5

12. Bardot ES, Hadjantonakis AK. Mouse gastrulation: Coordination of tissue patterning, specification and diversification of cell fate. Mech Dev. Sep 2020;163:103617. doi:10.1016/j.mod.2020.103617

13. Morgani SM, Hadjantonakis AK. Signaling regulation during gastrulation: Insights from mouse embryos and in vitro systems. Curr Top Dev Biol. 2020;137:391–431. doi:10.1016/bs.ctdb.2019.11.011

14. Camus A, Perea-Gomez A, Moreau A, Collignon J. Absence of Nodal signaling promotes precocious neural differentiation in the mouse embryo. Dev Biol. Jul 2006;295(2):743–55. doi:10.1016/j.ydbio.2006.03.047

15. Mesnard D, Guzman-Ayala M, Constam DB. Nodal specifies embryonic visceral endoderm and sustains pluripotent cells in the epiblast before overt axial patterning. Development. Jul 2006;133(13):2497–505. doi:10.1242/dev.02413

16. Xiao L, Yuan X, Sharkis SJ. Activin A maintains self-renewal and regulates fibroblast growth factor, Wnt, and bone morphogenic protein pathways in human embryonic stem cells. Stem Cells. Jun 2006;24(6):1476–86. doi:10.1634/stemcells.2005-0299

17. Vallier L, Mendjan S, Brown S, et al. Activin/Nodal signalling maintains pluripotency by controlling Nanog expression. Development. Apr 2009;136(8):1339–49. doi:10.1242/dev.033951

18. Liu P, Wakamiya M, Shea MJ, Albrecht U, Behringer RR, Bradley A. Requirement for Wnt3 in vertebrate axis formation. Nat Genet. Aug 1999;22(4):361–5. doi:10.1038/11932

19. Yoon Y, Huang T, Tortelote GG, et al. Extra-embryonic Wnt3 regulates the establishment of the primitive streak in mice. Dev Biol. Jul 1 2015;403(1):80–8. doi:10.1016/j.ydbio.2015.04.008

20. Funa NS, Schachter KA, Lerdrup M, et al. β-Catenin Regulates Primitive Streak Induction through Collaborative Interactions with SMAD2/SMAD3 and OCT4. Cell Stem Cell. Jun 4 2015;16(6):639–52. doi:10.1016/j.stem.2015.03.008

21. Estarás C, Benner C, Jones KA. SMADs and YAP compete to control elongation of β- catenin:LEF-1-recruited RNAPII during hESC differentiation. Mol Cell. Jun 2015;58(5):780–93. doi:10.1016/j.molcel.2015.04.001

22. Gadue P, Huber TL, Paddison PJ, Keller GM. Wnt and TGF-beta signaling are required for the induction of an in vitro model of primitive streak formation using embryonic stem cells. Proc Natl Acad Sci U S A. Nov 07 2006;103(45):16806–11. doi:10.1073/pnas.0603916103

23. Robertson EJ. Dose-dependent Nodal/Smad signals pattern the early mouse embryo. Semin Cell Dev Biol. Aug 2014;32:73–9. doi:10.1016/j.semcdb.2014.03.028

24. Arkell RM, Fossat N, Tam PP. Wnt signalling in mouse gastrulation and anterior development: new players in the pathway and signal output. Curr Opin Genet Dev. Aug 2013;23(4):454–60. doi:10.1016/j.gde.2013.03.001

25. Stronati E, Giraldez S, Huang L, et al. YAP1 regulates the self-organized fate patterning of hESC-derived gastruloids. Stem Cell Reports. Jan 03 2022;doi:10.1016/j.stemcr.2021.12.012

26. Hsu HT, Estarás C, Huang L, Jones KA. Specifying the Anterior Primitive Streak by Modulating YAP1 Levels in Human Pluripotent Stem Cells. Stem Cell Reports. 12 2018;11(6):1357–1364. doi:10.1016/j.stemcr.2018.10.013

27. Beyer TA, Weiss A, Khomchuk Y, et al. Switch enhancers interpret TGF-β and Hippo signaling to control cell fate in human embryonic stem cells. Cell Rep. Dec 2013;5(6):1611–24. doi:10.1016/j.celrep.2013.11.021

28. Abraham E, Kostina A, Volmert B, et al. A retinoic acid:YAP1 signaling axis controls atrial lineage commitment. Cell Rep. May 07 2025;44(5):115687. doi:10.1016/j.celrep.2025.115687

29. Hayashi S, Lewis P, Pevny L, McMahon AP. Efficient gene modulation in mouse epiblast using a Sox2Cre transgenic mouse strain. Mech Dev. Dec 2002;119 Suppl 1:S97–S101. doi:10.1016/s0925-4773(03)00099-6

30. Abraham E, Zubillaga M, Roule T, Stronati E, Akizu N, Estaras C. Single-Cell RNA Sequencing of Mutant Whole Mouse Embryos: From the Epiblast to the End of Gastrulation. J Vis Exp. Jun 14 2024;(208)doi:10.3791/66866

31. Argelaguet R, Clark SJ, Mohammed H, et al. Multi-omics profiling of mouse gastrulation at single-cell resolution. Nature. 12 2019;576(7787):487-491. doi:10.1038/s41586-019-1825-8

32. Pijuan-Sala B, Griffiths JA, Guibentif C, et al. A single-cell molecular map of mouse gastrulation and early organogenesis. Nature. 02 2019;566(7745):490-495. doi:10.1038/s41586-019-0933-9

33. Mittnenzweig M, Mayshar Y, Cheng S, et al. A single-embryo, single-cell time-resolved model for mouse gastrulation. Cell. May 27 2021;184(11):2825–2842.e22. doi:10.1016/j.cell.2021.04.004

34. Hirate Y, Hirahara S, Inoue K, et al. Polarity-dependent distribution of angiomotin localizes Hippo signaling in preimplantation embryos. Curr Biol. Jul 08 2013;23(13):1181–94. doi:10.1016/j.cub.2013.05.014

35. Blakely WJ, Hatterschide J, White EA. HPV18 E7 inhibits LATS1 kinase and activates YAP1 by degrading PTPN14. mBio. Oct 16 2024;15(10):e0181124. doi:10.1128/mbio.01811-24

36. Hermann A, Wu G, Nedvetsky PI, et al. The Hippo pathway component Wwc2 is a key regulator of embryonic development and angiogenesis in mice. Cell Death Dis. Jan 22 2021;12(1):117. doi:10.1038/s41419-021-03409-0

37. Estarás C, Hsu HT, Huang L, Jones KA. YAP repression of the WNT3 gene controls hESC differentiation along the cardiac mesoderm lineage. Genes Dev. 11 2017;31(22):2250–2263. doi:10.1101/gad.307512.117

38. Xiao F, Liao B, Hu J, et al. JMJD1C Ensures Mouse Embryonic Stem Cell Self-Renewal and Somatic Cell Reprogramming through Controlling MicroRNA Expression. Stem Cell Reports. Sep 12 2017;9(3):927–942. doi:10.1016/j.stemcr.2017.07.013

39. Dai HQ, Wang BA, Yang L, et al. TET-mediated DNA demethylation controls gastrulation by regulating Lefty-Nodal signalling. Nature. 10 2016;538(7626):528-532. doi:10.1038/nature20095

40. Dixon G, Pan H, Yang D, et al. QSER1 protects DNA methylation valleys from de novo methylation. Science. Apr 09 2021;372(6538)doi:10.1126/science.abd0875

41. Conlon FL, Lyons KM, Takaesu N, et al. A primary requirement for nodal in the formation and maintenance of the primitive streak in the mouse. Development. Jul 1994;120(7):1919–28.

42. Sun X, Meyers EN, Lewandoski M, Martin GR. Targeted disruption of Fgf8 causes failure of cell migration in the gastrulating mouse embryo. Genes Dev. Jul 15 1999;13(14):1834–46. doi:10.1101/gad.13.14.1834

43. Guo Q, Li JY. Distinct functions of the major Fgf8 spliceform, Fgf8b, before and during mouse gastrulation. Development. Jun 2007;134(12):2251-60. doi:10.1242/dev.004929

44. Hernández-Martínez R, Nowotschin S, Harland LTG, et al. Axin1 and Axin2 regulate the WNT-signaling landscape to promote distinct mesoderm programs. bioRxiv. Sep 11 2024;doi:10.1101/2024.09.11.612342

45. Wang Z, Oron E, Nelson B, Razis S, Ivanova N. Distinct lineage specification roles for NANOG, OCT4, and SOX2 in human embryonic stem cells. Cell Stem Cell. Apr 2012;10(4):440-54. doi:10.1016/j.stem.2012.02.016

46. Xu RH, Sampsell-Barron TL, Gu F, et al. NANOG is a direct target of TGFbeta/activin-mediated SMAD signaling in human ESCs. Cell Stem Cell. Aug 2008;3(2):196–206. doi:10.1016/j.stem.2008.07.001

47. Yu MS, Spiering S, Colas AR. Generation of First Heart Field-like Cardiac Progenitors and Ventricular-like Cardiomyocytes from Human Pluripotent Stem Cells. J Vis Exp. 06 2018;(136)doi:10.3791/57688

48. Malaguti M, Migueles RP, Blin G, Lin CY, Lowell S. Id1 Stabilizes Epiblast Identity by Sensing Delays in Nodal Activation and Adjusting the Timing of Differentiation. Dev Cell. Aug 19 2019;50(4):462–477.e5. doi:10.1016/j.devcel.2019.05.032

49. Cunningham TJ, Yu MS, McKeithan WL, et al. Id genes are essential for early heart formation. Genes Dev. Jul 1 2017;31(13):1325–1338. doi:10.1101/gad.300400.117

50. Martyn I, Brivanlou AH, Siggia ED. A wave of WNT signaling balanced by secreted inhibitors controls primitive streak formation in micropattern colonies of human embryonic stem cells. Development. 03 25 2019;146(6)doi:10.1242/dev.172791

51. Stott D, Kispert A, Herrmann BG. Rescue of the tail defect of Brachyury mice. Genes Dev. Feb 1993;7(2):197–203. doi:10.1101/gad.7.2.197

52. Rivera-Pérez JA, Magnuson T. Primitive streak formation in mice is preceded by localized activation of Brachyury and Wnt3. Dev Biol. Dec 15 2005;288(2):363–71. doi:10.1016/j.ydbio.2005.09.012

53. Bulger EA, Muncie-Vasic I, Libby ARG, McDevitt TC, Bruneau BG. TBXT dose sensitivity and the decoupling of nascent mesoderm specification from EMT progression in 2D human gastruloids. Development. Mar 15 2024;151(6)doi:10.1242/dev.202516

54. Arnold SJ, Hofmann UK, Bikoff EK, Robertson EJ. Pivotal roles for eomesodermin during axis formation, epithelium-to-mesenchyme transition and endoderm specification in the mouse. Development. Feb 2008;135(3):501–11. doi:10.1242/dev.014357

55. Burtscher I, Lickert H. Foxa2 regulates polarity and epithelialization in the endoderm germ layer of the mouse embryo. Development. Mar 2009;136(6):1029–38. doi:10.1242/dev.028415

56. Acampora D, Mazan S, Lallemand Y, et al. Forebrain and midbrain regions are deleted in Otx2-/- mutants due to a defective anterior neuroectoderm specification during gastrulation. Development. Oct 1995;121(10):3279–90. doi:10.1242/dev.121.10.3279

57. Stiles J, Jernigan TL. The basics of brain development. Neuropsychol Rev. Dec 2010;20(4):327–48. doi:10.1007/s11065-010-9148-4

58. LeBlanc L, Ramirez N, Kim J. Context-dependent roles of YAP/TAZ in stem cell fates and cancer. Cell Mol Life Sci. May 2021;78(9):4201–4219. doi:10.1007/s00018-021-03781-2

59. Handford CE, Junyent S, Jorgensen V, Zernicka-Goetz M. Topical section: embryonic models (2023) for Current Opinion in Genetics & Development. Curr Opin Genet Dev. Feb 2024;84:102134. doi:10.1016/j.gde.2023.102134

60. Rossant J, Tam PPL. Early human embryonic development: Blastocyst formation to gastrulation. Dev Cell. 01 24 2022;57(2):152-165. doi:10.1016/j.devcel.2021.12.022

61. Brons IG, Smithers LE, Trotter MW, et al. Derivation of pluripotent epiblast stem cells from mammalian embryos. Nature. Jul 12 2007;448(7150):191-5. doi:10.1038/nature05950

62. Warrier S, Van der Jeught M, Duggal G, et al. Direct comparison of distinct naive pluripotent states in human embryonic stem cells. Nat Commun. Apr 21 2017;8:15055. doi:10.1038/ncomms15055

63. Kim DI, Jensen SC, Noble KA, et al. An improved smaller biotin ligase for BioID proximity labeling. Mol Biol Cell. Apr 2016;27(8):1188–96. doi:10.1091/mbc.E15-12-0844

64. Zanconato F, Battilana G, Forcato M, et al. Transcriptional addiction in cancer cells is mediated by YAP/TAZ through BRD4. Nat Med. Oct 2018;24(10):1599–1610. doi:10.1038/s41591-018-0158-8

65. Chang L, Azzolin L, Di Biagio D, et al. The SWI/SNF complex is a mechanoregulated inhibitor of YAP and TAZ. Nature. Nov 2018;563(7730):265-269. doi:10.1038/s41586-018-0658-1

66. Fasciani A, D’Annunzio S, Poli V, et al. MLL4-associated condensates counterbalance Polycomb-mediated nuclear mechanical stress in Kabuki syndrome. Nat Genet. Dec 2020;52(12):1397–1411. doi:10.1038/s41588-020-00724-8

67. Liu W, Huang Y, Wang D, et al. MPDZ as a novel epigenetic silenced tumor suppressor inhibits growth and progression of lung cancer through the Hippo-YAP pathway. Oncogene. Jul 2021;40(26):4468–4485. doi:10.1038/s41388-021-01857-8

68. Barry ER, Morikawa T, Butler BL, et al. Restriction of intestinal stem cell expansion and the regenerative response by YAP. Nature. Jan 2013;493(7430):106-10. doi:10.1038/nature11693

69. Kloet SL, Karemaker ID, van Voorthuijsen L, et al. NuRD-interacting protein ZFP296 regulates genome-wide NuRD localization and differentiation of mouse embryonic stem cells. Nat Commun. Nov 02 2018;9(1):4588. doi:10.1038/s41467-018-07063-7

70. Kueng S, Hegemann B, Peters BH, et al. Wapl controls the dynamic association of cohesin with chromatin. Cell. Dec 1 2006;127(5):955–67. doi:10.1016/j.cell.2006.09.040

71. Wu MY, Ramel MC, Howell M, Hill CS. SNW1 is a critical regulator of spatial BMP activity, neural plate border formation, and neural crest specification in vertebrate embryos. PLoS Biol. Feb 15 2011;9(2):e1000593. doi:10.1371/journal.pbio.1000593

72. Zhao X, Fang K, Liu X, et al. QSER1 preserves the suppressive status of the pro-apoptotic genes to prevent apoptosis. Cell Death Differ. Mar 2023;30(3):779–793. doi:10.1038/s41418-022-01085-x

73. Zhu C, Li L, Zhang Z, et al. A Non-canonical Role of YAP/TEAD Is Required for Activation of Estrogen-Regulated Enhancers in Breast Cancer. Mol Cell. 08 2019;75(4):791–806.e8. doi:10.1016/j.molcel.2019.06.010

74. Minde DP, Ramakrishna M, Lilley KS. Biotin proximity tagging favours unfolded proteins and enables the study of intrinsically disordered regions. Commun Biol. Jan 22 2020;3(1):38. doi:10.1038/s42003-020-0758-y

75. Mesrouze Y, Bokhovchuk F, Izaac A, et al. Adaptation of the bound intrinsically disordered protein YAP to mutations at the YAP:TEAD interface. Protein Sci. Oct 2018;27(10):1810–1820. doi:10.1002/pro.3493

76. Zhao B, Ye X, Yu J, et al. TEAD mediates YAP-dependent gene induction and growth control. Genes Dev. Jul 2008;22(14):1962–71. doi:10.1101/gad.1664408

77. Hao Y, Chun A, Cheung K, Rashidi B, Yang X. Tumor suppressor LATS1 is a negative regulator of oncogene YAP. J Biol Chem. Feb 2008;283(9):5496–509. doi:10.1074/jbc.M709037200

78. Zhao B, Li L, Lu Q, et al. Angiomotin is a novel Hippo pathway component that inhibits YAP oncoprotein. Genes Dev. Jan 1 2011;25(1):51–63. doi:10.1101/gad.2000111

79. Abramson J, Adler J, Dunger J, et al. Accurate structure prediction of biomolecular interactions with AlphaFold 3. Nature. Jun 2024;630(8016):493-500. doi:10.1038/s41586-024-07487-w

80. Rai A, Oprisko A, Campos J, et al. bMERB domains are bivalent Rab8 family effectors evolved by gene duplication. Elife. Aug 23 2016;5doi:10.7554/eLife.18675

81. Rada-Iglesias A, Bajpai R, Swigut T, Brugmann SA, Flynn RA, Wysocka J. A unique chromatin signature uncovers early developmental enhancers in humans. Nature. Feb 2011;470(7333):279-83. doi:10.1038/nature09692

82. Kagey MH, Newman JJ, Bilodeau S, et al. Mediator and cohesin connect gene expression and chromatin architecture. Nature. Sep 2010;467(7314):430-5. doi:10.1038/nature09380

83. Kanehisa M, Sato Y, Kawashima M, Furumichi M, Tanabe M. KEGG as a reference resource for gene and protein annotation. Nucleic Acids Res. Jan 4 2016;44(D1):D457–62. doi:10.1093/nar/gkv1070

84. Abdel Mouti M, Pauklin S. TGFB1/INHBA Homodimer/Nodal-SMAD2/3 Signaling Network: A Pivotal Molecular Target in PDAC Treatment. Mol Ther. Mar 3 2021;29(3):920–936. doi:10.1016/j.ymthe.2021.01.002

85. Namwanje M, Brown CW. Activins and Inhibins: Roles in Development, Physiology, and Disease. Cold Spring Harb Perspect Biol. Jul 1 2016;8(7)doi:10.1101/cshperspect.a021881

86. Powers SE, Taniguchi K, Yen W, et al. Tgif1 and Tgif2 regulate Nodal signaling and are required for gastrulation. Development. Jan 2010;137(2):249–59. doi:10.1242/dev.040782

87. Zinski J, Tajer B, Mullins MC. TGF-β Family Signaling in Early Vertebrate Development. Cold Spring Harb Perspect Biol. 06 2018;10(6)doi:10.1101/cshperspect.a033274

88. Lian X, Zhang J, Azarin SM, et al. Directed cardiomyocyte differentiation from human pluripotent stem cells by modulating Wnt/β-catenin signaling under fully defined conditions. Nat Protoc. Jan 2013;8(1):162–75. doi:10.1038/nprot.2012.150

89. He W, Lu Q, Sherchan P, et al. Activation of Frizzled-7 attenuates blood-brain barrier disruption through Dvl/β-catenin/WISP1 signaling pathway after intracerebral hemorrhage in mice. Fluids Barriers CNS. Sep 26 2021;18(1):44. doi:10.1186/s12987-021-00278-9

90. Scheibner K, Schirge S, Burtscher I, et al. Epithelial cell plasticity drives endoderm formation during gastrulation. Nat Cell Biol. Jul 2021;23(7):692–703. doi:10.1038/s41556-021-00694-x

91. Hagenbeek TJ, Zbieg JR, Hafner M, et al. An allosteric pan-TEAD inhibitor blocks oncogenic YAP/TAZ signaling and overcomes KRAS G12C inhibitor resistance. Nat Cancer. Jun 2023;4(6):812–828. doi:10.1038/s43018-023-00577-0

92. Dunbrack RL, Jr. Rēs ipSAE loquunt: What’s wrong with AlphaFold’s ipTM score and how to fix it. *bioRxiv*. Feb 14 2025;doi:10.1101/2025.02.10.637595

93. Vincent SD, Dunn NR, Hayashi S, Norris DP, Robertson EJ. Cell fate decisions within the mouse organizer are governed by graded Nodal signals. Genes Dev. Jul 2003;17(13):1646–62. doi:10.1101/gad.1100503

94. Deglincerti A, Etoc F, Guerra MC, et al. Self-organization of human embryonic stem cells on micropatterns. Nat Protoc. Nov 2016;11(11):2223–2232. doi:10.1038/nprot.2016.131

95. Liu L, Nemashkalo A, Rezende L, et al. Nodal is a short-range morphogen with activity that spreads through a relay mechanism in human gastruloids. Nat Commun. Jan 25 2022;13(1):497. doi:10.1038/s41467-022-28149-3

96. Tang Z, Ma Q, Wang L, et al. A brief review: some compounds targeting YAP against malignancies. Future Oncol. May 2019;15(13):1535–1543. doi:10.2217/fon-2019-0035

97. Rito T, Libby ARG, Demuth M, Domart MC, Cornwall-Scoones J, Briscoe J. Timely TGFβ signalling inhibition induces notochord. Nature. Dec 18 2024;doi:10.1038/s41586-024-08332-w

98. Wotton D, Lo RS, Swaby LA, Massagué J. Multiple modes of repression by the Smad transcriptional corepressor TGIF. J Biol Chem. Dec 24 1999;274(52):37105–10. doi:10.1074/jbc.274.52.37105

99. Gripp KW, Wotton D, Edwards MC, et al. Mutations in TGIF cause holoprosencephaly and link NODAL signalling to human neural axis determination. Nat Genet. Jun 2000;25(2):205–8. doi:10.1038/76074

100. Iratni R, Yan YT, Chen C, et al. Inhibition of excess nodal signaling during mouse gastrulation by the transcriptional corepressor DRAP1. Science. Dec 6 2002;298(5600):1996-9. doi:10.1126/science.1073405

101. Meno C, Gritsman K, Ohishi S, et al. Mouse Lefty2 and zebrafish antivin are feedback inhibitors of nodal signaling during vertebrate gastrulation. Mol Cell. Sep 1999;4(3):287–98. doi:10.1016/s1097-2765(00)80331-7

102. Perea-Gomez A, Vella FD, Shawlot W, et al. Nodal antagonists in the anterior visceral endoderm prevent the formation of multiple primitive streaks. Dev Cell. Nov 2002;3(5):745–56. doi:10.1016/s1534-5807(02)00321-0

103. Perea-Gomez A, Rhinn M, Ang SL. Role of the anterior visceral endoderm in restricting posterior signals in the mouse embryo. Int J Dev Biol. 2001;45(1):311–20.

104. Ferrie JJ, Karr JP, Tjian R, Darzacq X. "Structure"-function relationships in eukaryotic transcription factors: The role of intrinsically disordered regions in gene regulation. Mol Cell. Nov 3 2022;82(21):3970–3984. doi:10.1016/j.molcel.2022.09.021

105. He J, Huo X, Pei G, et al. Dual-role transcription factors stabilize intermediate expression levels. Cell. May 23 2024;187(11):2746–2766.e25. doi:10.1016/j.cell.2024.03.023

106. Galli GG, Carrara M, Yuan WC, et al. YAP Drives Growth by Controlling Transcriptional Pause Release from Dynamic Enhancers. Mol Cell. Oct 2015;60(2):328–37. doi:10.1016/j.molcel.2015.09.001

107. Hughes JR, Roberts N, McGowan S, et al. Analysis of hundreds of cis-regulatory landscapes at high resolution in a single, high-throughput experiment. Nat Genet. Feb 2014;46(2):205–12. doi:10.1038/ng.2871

108. Mumbach MR, Rubin AJ, Flynn RA, et al. HiChIP: efficient and sensitive analysis of protein-directed genome architecture. Nat Methods. Nov 2016;13(11):919–922. doi:10.1038/nmeth.3999

109. Siegl D, Kruchem M, Jansky S, et al. A PCR protocol to establish standards for routine mycoplasma testing that by design detects over ninety percent of all known mycoplasma species. iScience. May 19 2023;26(5):106724. doi:10.1016/j.isci.2023.106724

110. Hayashi S, Tenzen T, McMahon AP. Maternal inheritance of Cre activity in a Sox2Cre deleter strain. Genesis. Oct 2003;37(2):51–3. doi:10.1002/gene.10225

111. Ritchie ME, Phipson B, Wu D, et al. limma powers differential expression analyses for RNA-sequencing and microarray studies. Nucleic Acids Res. Apr 20 2015;43(7):e47. doi:10.1093/nar/gkv007

112. Jumper J, Evans R, Pritzel A, et al. Highly accurate protein structure prediction with AlphaFold. Nature. Aug 2021;596(7873):583-589. doi:10.1038/s41586-021-03819-2

113. Tunyasuvunakool K, Adler J, Wu Z, et al. Highly accurate protein structure prediction for the human proteome. Nature. Aug 2021;596(7873):590-596. doi:10.1038/s41586-021-03828-1

114. Kryshtafovych A, Schwede T, Topf M, Fidelis K, Moult J. Critical assessment of methods of protein structure prediction (CASP)-Round XIV. Proteins. Dec 2021;89(12):1607–1617. doi:10.1002/prot.26237

115. Huang SY, Zou X. A knowledge-based scoring function for protein-RNA interactions derived from a statistical mechanics-based iterative method. Nucleic Acids Res. Apr 2014;42(7):e55. doi:10.1093/nar/gku077

116. Huang SY, Zou X. An iterative knowledge-based scoring function for protein-protein recognition. Proteins. Aug 2008;72(2):557–79. doi:10.1002/prot.21949

117. Yan Y, Zhang D, Zhou P, Li B, Huang SY. HDOCK: a web server for protein-protein and protein-DNA/RNA docking based on a hybrid strategy. Nucleic Acids Res. Jul 3 2017;45(W1):W365–w373. doi:10.1093/nar/gkx407

118. Yan Y, Tao H, He J, Huang SY. The HDOCK server for integrated protein-protein docking. Nat Protoc. May 2020;15(5):1829–1852. doi:10.1038/s41596-020-0312-x

119. Yan Y, Wen Z, Wang X, Huang SY. Addressing recent docking challenges: A hybrid strategy to integrate template-based and free protein-protein docking. Proteins. Mar 2017;85(3):497–512. doi:10.1002/prot.25234

120. Wodak SJ, Vajda S, Lensink MF, Kozakov D, Bates PA. Critical Assessment of Methods for Predicting the 3D Structure of Proteins and Protein Complexes. Annu Rev Biophys. May 9 2023;52:183–206. doi:10.1146/annurev-biophys-102622-084607

121. Katchalski-Katzir E, Shariv I, Eisenstein M, Friesem AA, Aflalo C, Vakser IA. Molecular surface recognition: determination of geometric fit between proteins and their ligands by correlation techniques. Proc Natl Acad Sci U S A. Mar 15 1992;89(6):2195–9. doi:10.1073/pnas.89.6.2195

122. Case DA, Aktulga HM, Belfon K, et al. AmberTools. J Chem Inf Model. Oct 23 2023;63(20):6183-6191. doi:10.1021/acs.jcim.3c01153

123. Case DA, Cheatham TE, 3rd, Darden T, et al. The Amber biomolecular simulation programs. J Comput Chem. Dec 2005;26(16):1668–88. doi:10.1002/jcc.20290

124. Anandakrishnan R, Aguilar B, Onufriev AV. H++ 3.0: automating pK prediction and the preparation of biomolecular structures for atomistic molecular modeling and simulations. Nucleic Acids Res. Jul 2012;40(Web Server issue):W537-41. doi:10.1093/nar/gks375

125. Tian C, Kasavajhala K, Belfon KAA, et al. ff19SB: Amino-Acid-Specific Protein Backbone Parameters Trained against Quantum Mechanics Energy Surfaces in Solution. J Chem Theory Comput. Jan 14 2020;16(1):528–552. doi:10.1021/acs.jctc.9b00591

126. Izadi S, Anandakrishnan R, Onufriev AV. Building Water Models: A Different Approach. J Phys Chem Lett. Nov 6 2014;5(21):3863–3871. doi:10.1021/jz501780a

127. Li P, Song LF, Merz KM, Jr. Systematic Parameterization of Monovalent Ions Employing the Nonbonded Model. J Chem Theory Comput. Apr 14 2015;11(4):1645–57. doi:10.1021/ct500918t

128. Abraham E, Volmert B, Roule T, et al. A Retinoic Acid:YAP1 signaling axis controls atrial lineage commitment. bioRxiv. Jul 12 2024;doi:10.1101/2024.07.11.602981

129. Meng EC, Goddard TD, Pettersen EF, et al. UCSF ChimeraX: Tools for structure building and analysis. Protein Sci. Nov 2023;32(11):e4792. doi:10.1002/pro.4792

130. Mirdita M, Schütze K, Moriwaki Y, Heo L, Ovchinnikov S, Steinegger M. ColabFold: making protein folding accessible to all. Nat Methods. Jun 2022;19(6):679–682. doi:10.1038/s41592-022-01488-1

