## Supplementary figures and images for "YAP1 and QSER1 are Key Modulators of Embryonic Signaling Pathways in the Mammalian Epiblast"

### Figure EV1

#### Expanded View 1

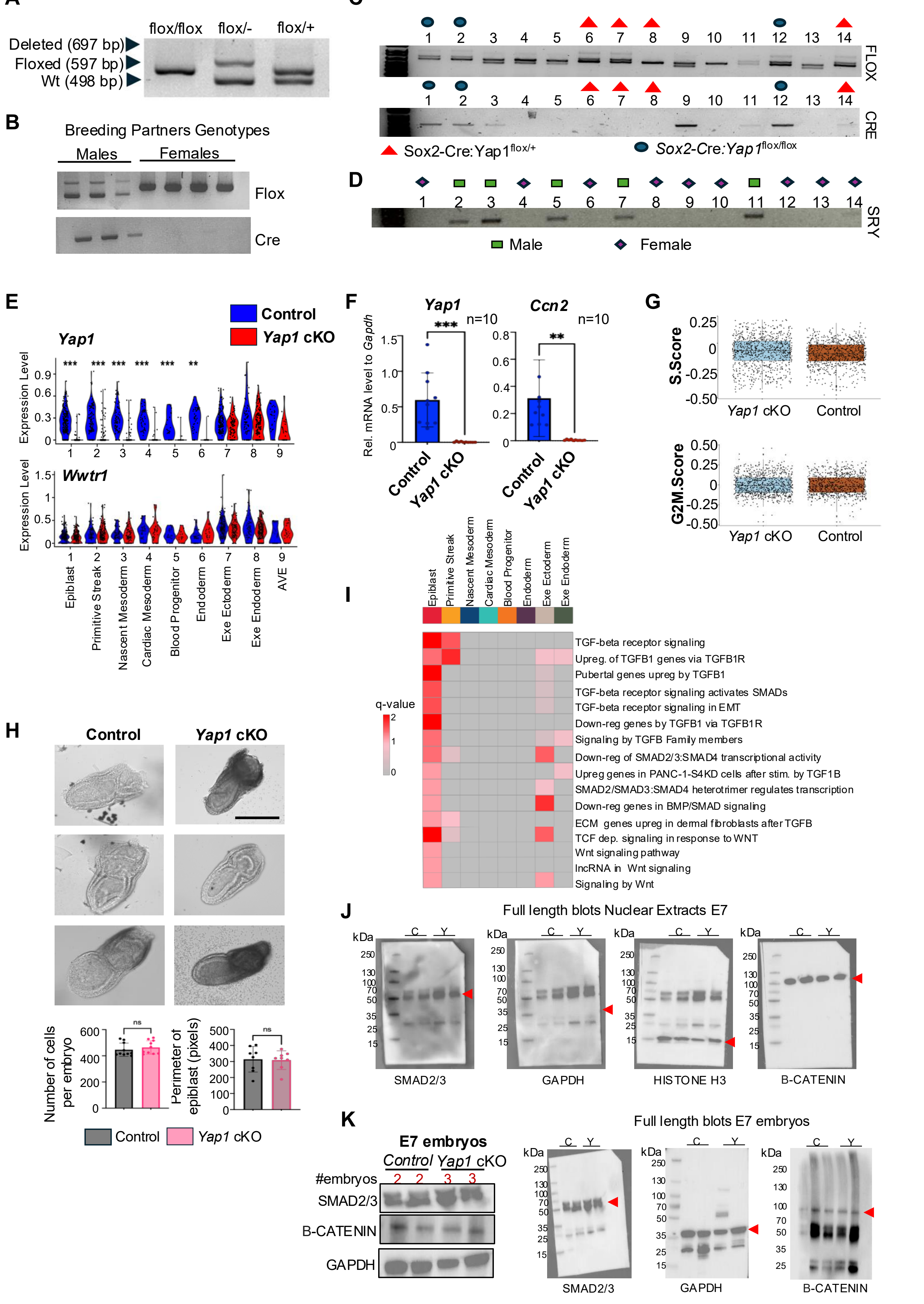

### Figure EV4

#### Expanded View 4

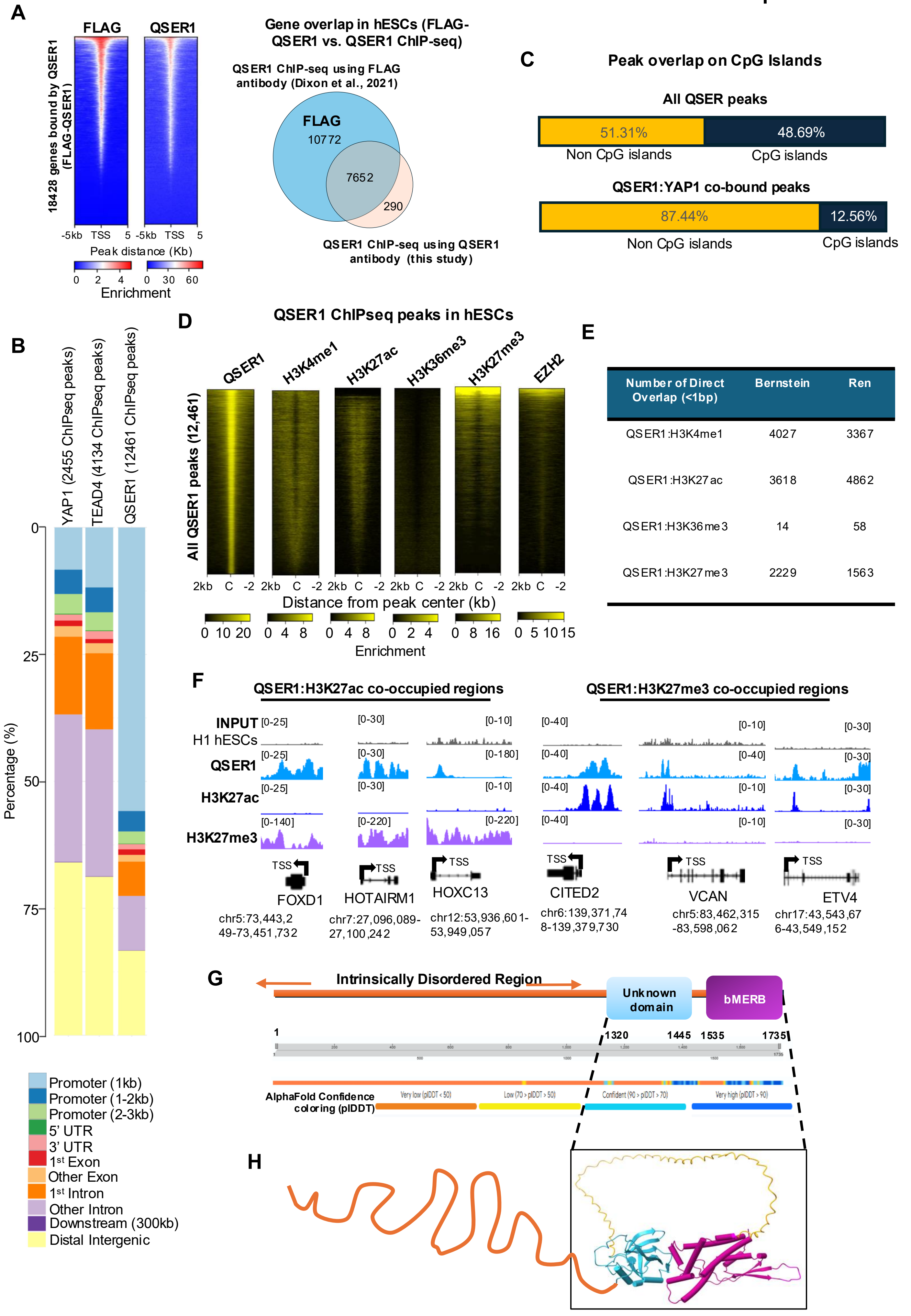


| Number of Direct Overlap (<1bp) | Bernstein | Ren |
|---|---|---|
| QSER1:H3K4me1 | 4027 | 3367 |
| QSER1:H3K27ac | 3618 | 4862 |
| QSER1:H3K36me3 | 14 | 58 |
| QSER1:H3K27me3 | 2229 | 1563 |

### Figure EV5

**Expanded View 5**

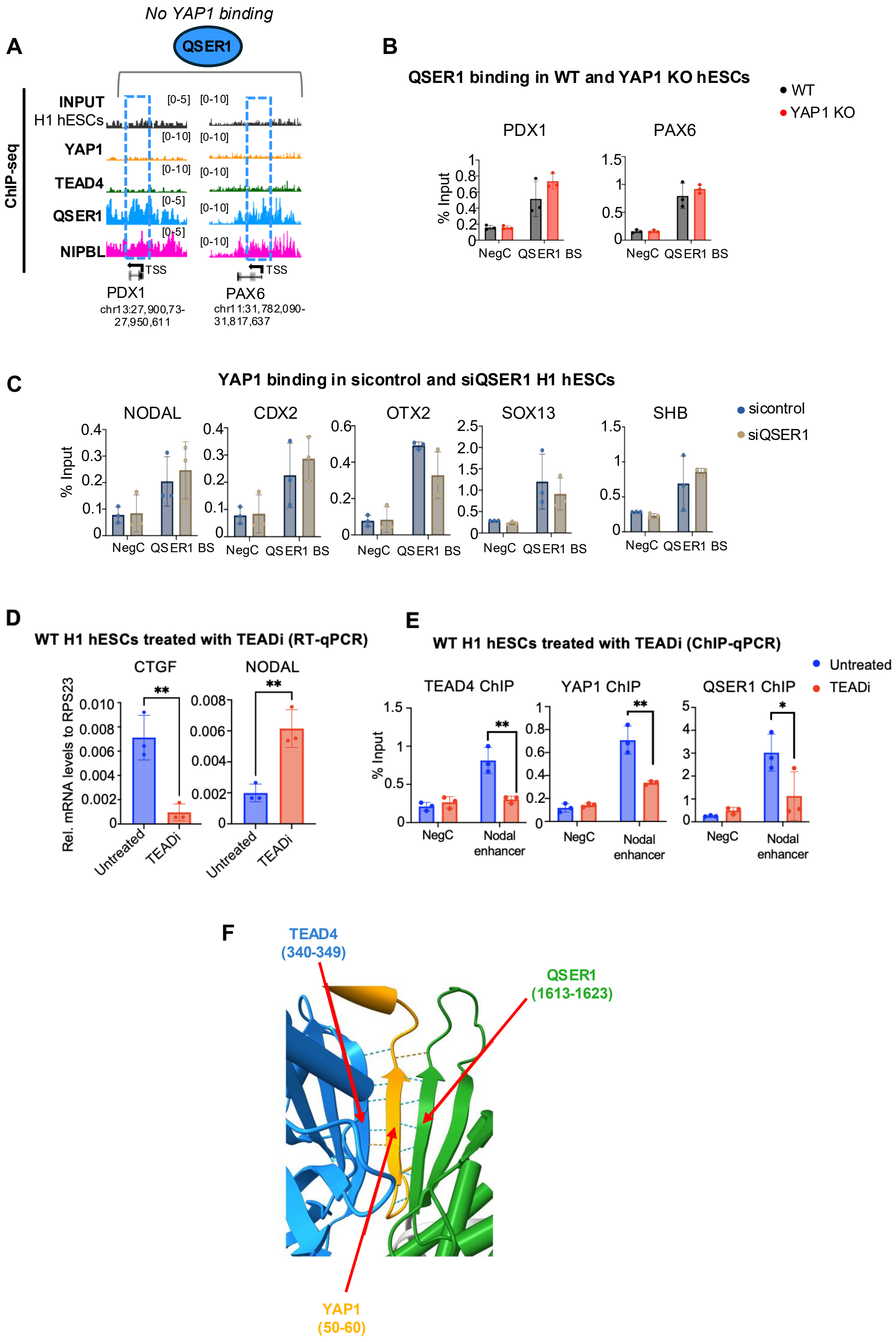

### Figure EV6

#### Expanded View 6

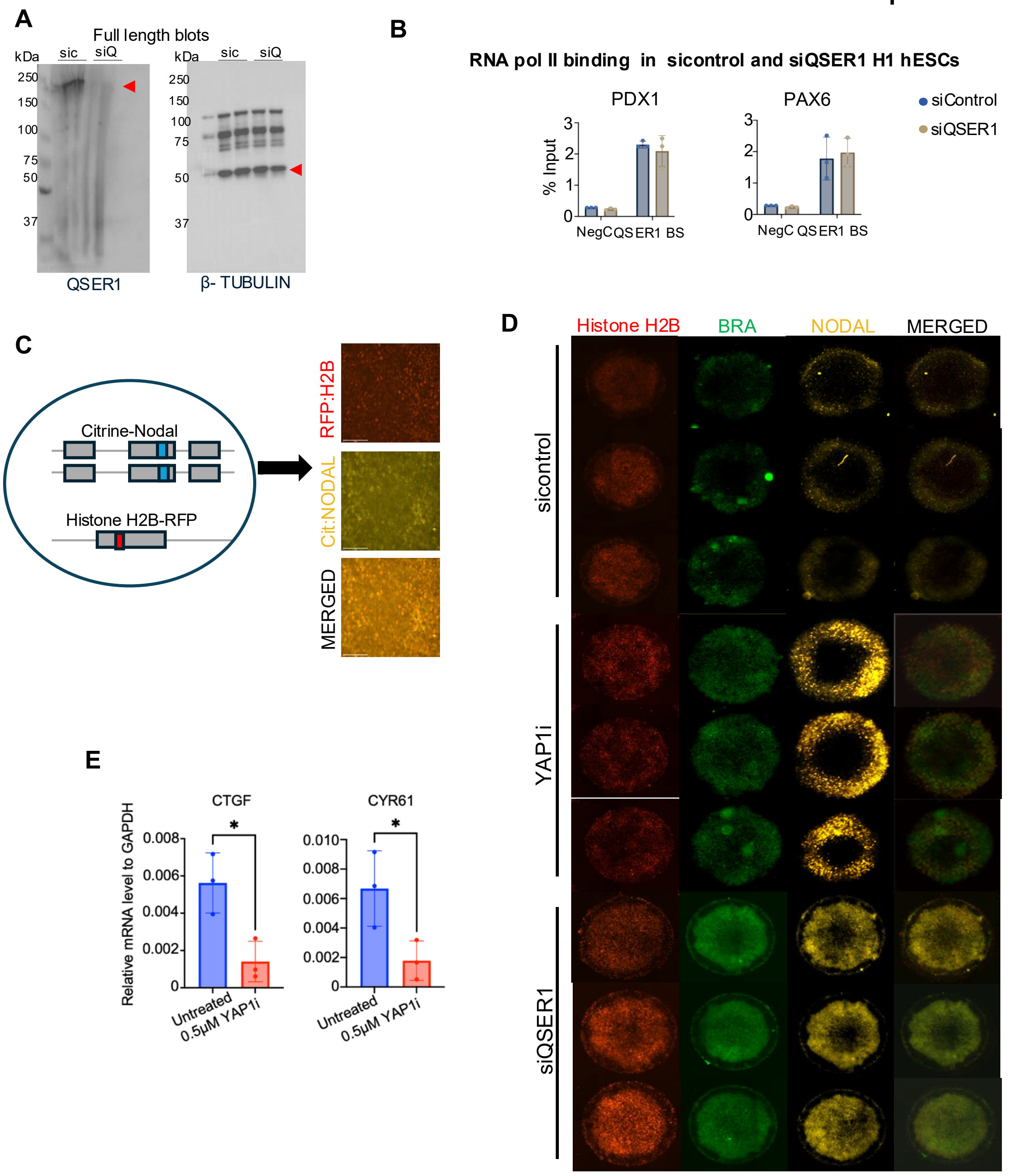
