## Supplementary material for "YAP1 and QSER1 are Key Modulators of Embryonic Signaling Pathways in the Mammalian Epiblast": Figure EV2

### Expanded View 2

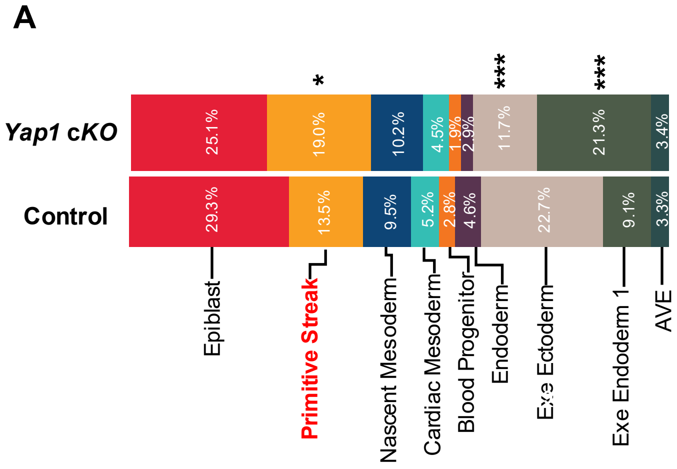


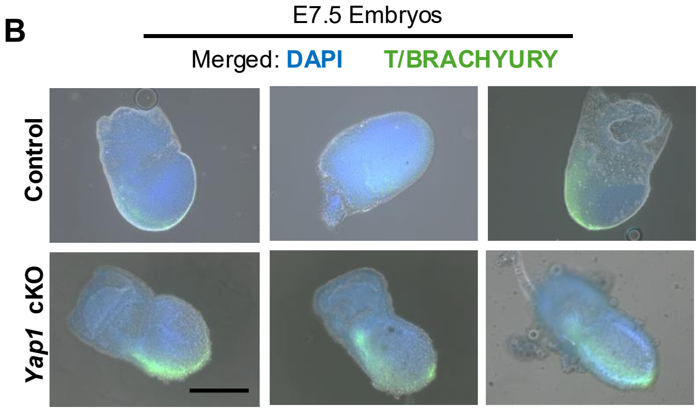


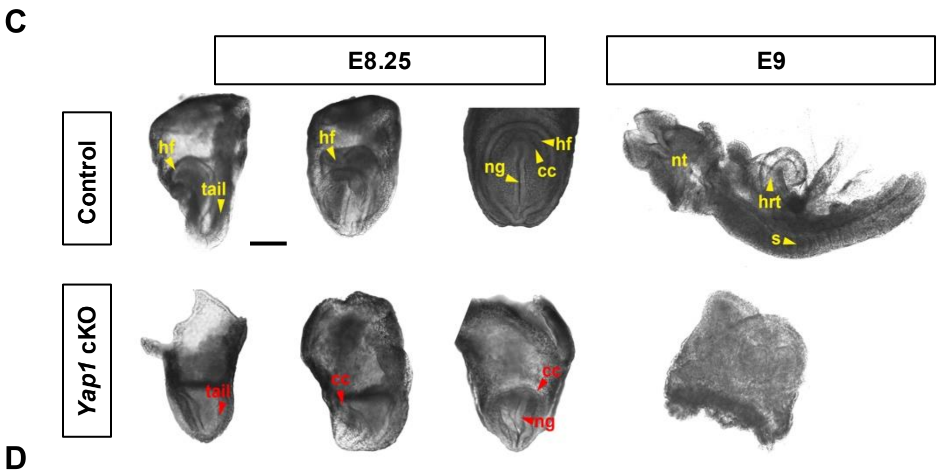


D

| E8.25 Phenotyping | Control | *Yap1* cKO |
|---|---|---|
| Head folds (hf) | Present (n=15) | Not visible in any of the embryos (n=15) |
| Cardiac crescent (cc) | Present (n=15) | Visible in 2/3 of the embryos (n=15) |
| A-P axis | Headfold and tail clearly defined along an A-P axis (n=15) | A-P axis short or not distinguishable (n=15) |

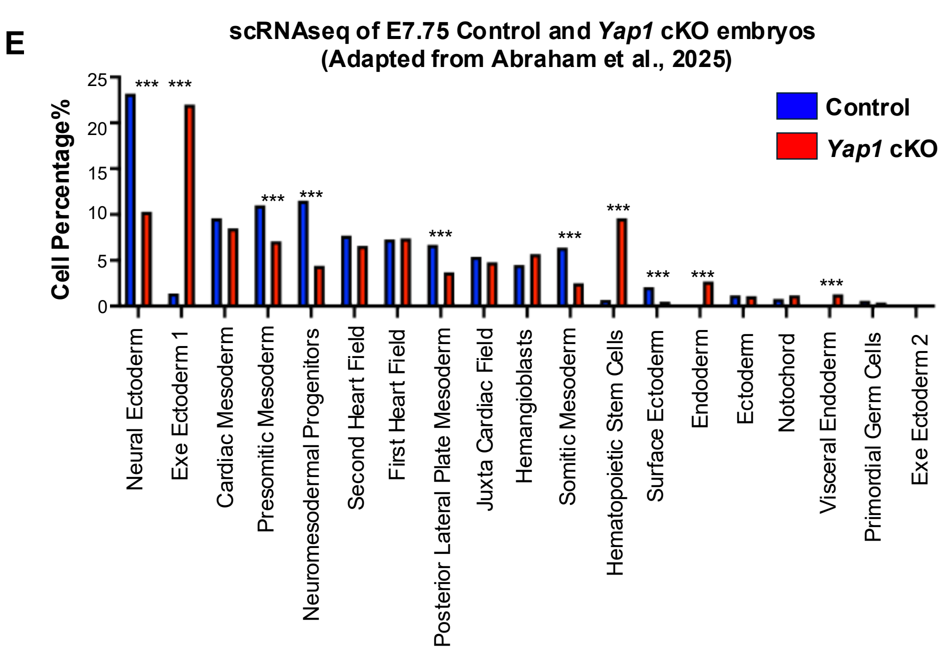


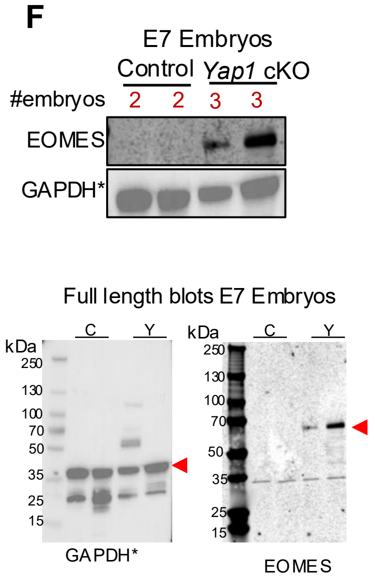


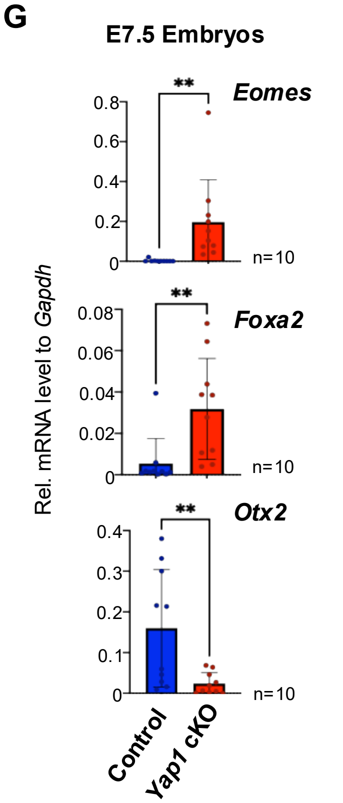
