## Supplementary material for "YAP1 and QSER1 are Key Modulators of Embryonic Signaling Pathways in the Mammalian Epiblast": Figure EV3

### Expanded View 3

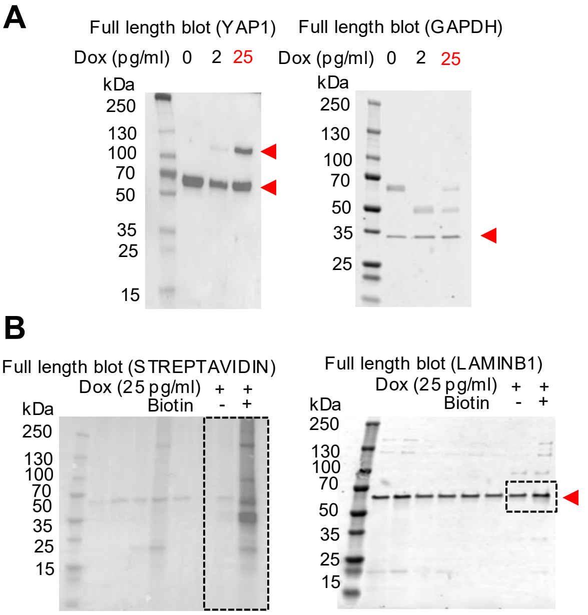


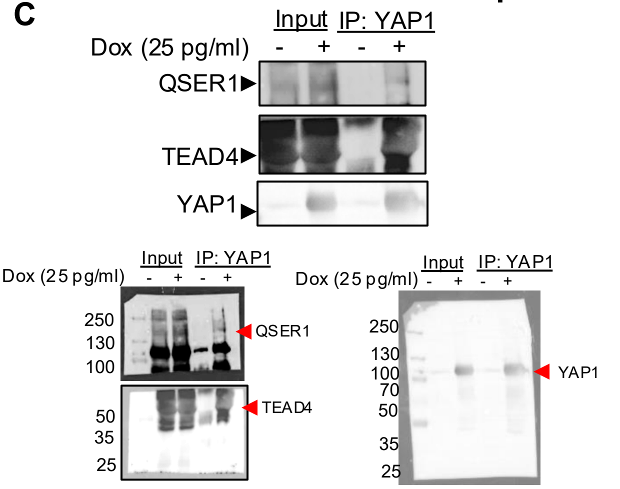


E

Raw MS/MS spectral counts

| Experimental Conditions | Plus Dox Plus Biotin | Plus Dox Minus Biotin | Minus Dox Plus Biotin | Minus Dox Minus Biotin |
|---|---|---|---|---|
| Biological Reps. | Rep1 / Rep2 / Rep3 | Rep1 / Rep2 / Rep3 | Rep1 / Rep2 / Rep3 | Rep1 / Rep2 / Lost |
| YAP1 | 66 / 160 / 45 | 43 / 59 / 140 | 0 / 0 / 2 | 0 / 0 / Lost |
| TEAD1 | 0 / 4 / 1 | 0 / 0 / 1 | 0 / 0 / 0 | 0 / 0 / Lost |
| TEAD2 | 0 / 1 / 0 | 0 / 0 / 0 | 0 / 0 / 0 | 0 / 0 / Lost |
| TEAD4 | 0 / 3 / 0 | 0 / 0 / 0 | 2 / 1 / 0 | 1 / 0 / Lost |
| QSER1 | 9/26/12 | 6/2/16 | 3 / 3 / 0 | 3 / 2 / Lost |
| AMOT | 11 / 45 / 0 | 0 / 0 / 10 | 0 / 0 / 0 | 0 / 0 / Lost |

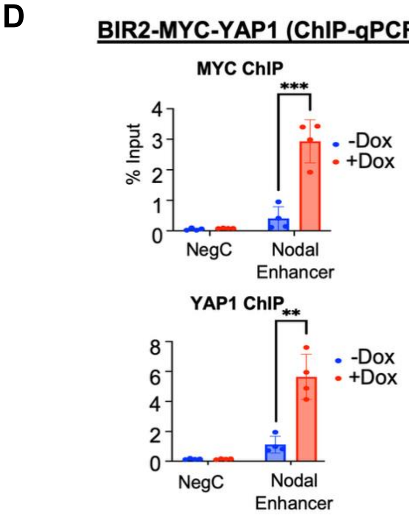


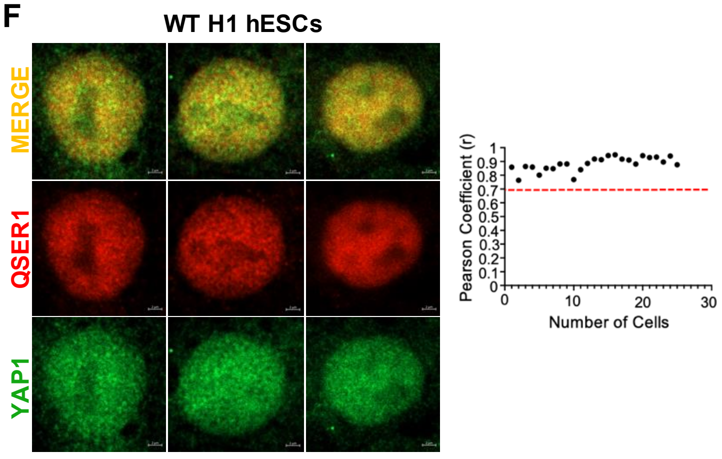


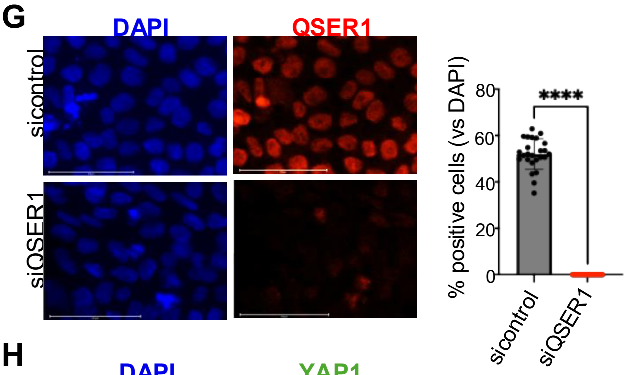


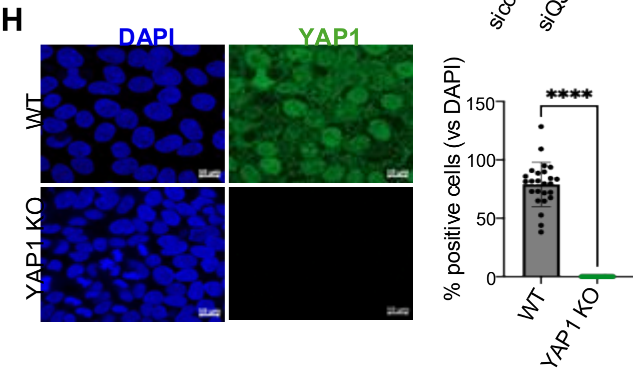


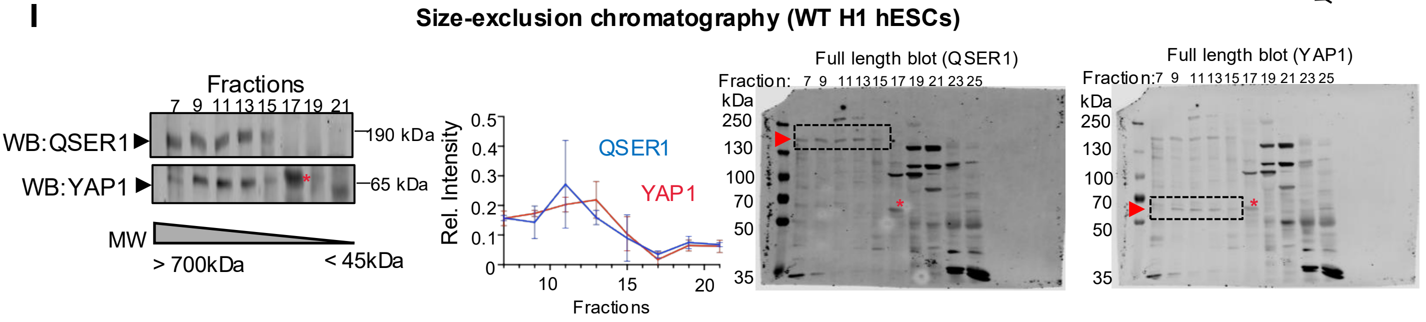
